# Transcriptomic Illuminations: How light intensity reshapes the *Chlamydomonas reinhardtii* cell cycle

**DOI:** 10.64898/2026.07.26.740802

**Authors:** Firas Louis, P.V. Sijil, Bipasha Bhattarcharjee, Martin Mora-Garcia, Rabinder Singh, Kateřina Bišová

**Affiliations:** Laboratory of Cell Cycles of Algae, Centre Algatech, Institute of Microbiology of the Czech Academy of Sciences, 237, Trebon 379 01, Czech Republic; Department of Cell biology and Molecular Genetics, University of Maryland, College Park MD 20742, USA

**Keywords:** light, commitment point, cell cycle, cell division, physiological responses, *Chlamydomonas*

## Abstract

The unicellular algae cell cycle can be divided into several phases, including the commitment point (CP), a point-of-no-return where the cell decides to divide, presumably based on reaching a critical cell size. Light plays a crucial role in the fitness of photosynthetic algal cells, affecting both CP timing and the number of daughter cells produced. So far, only few genes involved in CP have been described, and the presumed sizer and its signal(s) remain unidentified. Using synchronized cells and varying light intensities as a proxy, we explored the effects of light intensity in *Chlamydomonas reinhardtii* and observed both physiological and transcriptional changes occurring before and after CP under low light (LL) (100 µmol m⁻² s⁻¹) and optimal light (OL) (500 µmol m⁻² s⁻¹) conditions. Although CP was delayed by approximately 6 hours in LL, resulting in smaller mother cells and fewer daughter cells, the cells divided at the same time in both conditions. Overall, nucleic acid, protein, and energy reserve levels were lower in LL, with almost no starch produced. RNA-seq analysis identified several core genes shared between both conditions, with 201 genes expressed only in pre-CP1, 161 genes specific to post-CP1, and 582 shared across different phases. In LL, RNA-seq analysis showed an increase in differentially expressed genes (DEGs) in pre-CP1 compared to post-CP1, with an emphasis on photosynthesis, RNA metabolism, and organelle production before commitment, and cell division-related pathways (microtubules, DNA recombination) after CP1. In OL, the number of DEGs increased in post-CP1 by approximately 41% compared to pre-CP1, with a strong emphasis on protein production throughout the cell cycle.

## 1. INTRODUCTION

Microalgae are microscopic photosynthetic organisms that live in diverse conditions and environments, ranging from soil and ponds to arctic regions ^1–3^. Their development is influenced by many factors, including temperature, trophic regime, pH, and CO2 supply^4–7^. Light plays a crucial role as an essential component of photosynthesis, the primary source of energy for phototrophic organisms. In algae, as in higher plants, photosynthesis consists of a series of reactions that occur in chloroplasts. These photosynthetic reactions are divided into light and “dark” phases. During the light phase, in the thylakoid membrane, two photosystems (PS) operate in parallel to generate a proton (H^+^) gradient. PSI absorbs photon energy through chlorophyll pigments to release electrons, leading to the production of NADPH from NADP+, while PSII splits water (H_2_O) into H^+^, dioxygen (O_2_), and electrons, which return to the chlorophyll through the electron transport chain. The proton gradient generated by these two systems drives the production of adenosine triphosphate (ATP) from adenosine diphosphate (ADP) and inorganic phosphate (Pi). This potential energy is then used in the “dark” phase (light is permissible but not required) to fix carbon into sugar or starch through the Calvin-Benson-Bassham cycle^8–10^. Given its critical role in phototrophic organisms, light properties such as regime, composition, and quality have been shown to affect algal growth and the production of macromolecules like starch or lipids^11–16^. The most important factor is light intensity, which correlates with the photosynthetic rate and influences metabolite generation^17^. The growth rate increases with light intensity up to a species-dependent maximum; for example, 500 µmol m^−2^ s^−1^ is optimal for *Chlamydomonas reinhardtii* (hereafter *Chlamydomonas*), and 250 µmol m^−2^ s^−1^ for *Desmodesmus quadricauda* and *Parachlorella kessleri,* although these are not the most commonly used conditions^11^. However, excessive light intensity is detrimental, resulting in the absorption of excess energy that damages the light apparatus in a process called photoinhibition. The photosynthetic machinery must then be repaired by replacing damaged proteins, which reduces its efficiency over time^18^.

In eukaryotes, the cell cycle (CC) consists of several phases: (1) the G1 phase, corresponding to growth; (2) the S phase, during which DNA is replicated; (3) the G2 phase, where the cell continues to grow and prepares for division; and (4) the M phase, which includes mitosis and cell division. CC control is conserved in eukaryotes, both in Opisthokonts (animals and fungi) and in Viridiplantae (land plants and algae), with many homologs of the same genes shared and performing the same functions, suggesting that it may have originated in the LECA (Last Eukaryotic Common Ancestor) itself^19–21^. The conserved genes include those involved in DNA replication machinery (initiation, polymerases, DNA binding, etc.) and repair, spindle formation (tubulins, kinesins), and several cyclin-dependent kinases (CDKs) and their regulators, which are responsible for the positive and negative feedback loops that regulate the cell cycle phases ^22,23^.

Most mother cells in the known living world divide into two daughter cells through mitosis followed by cytokinesis, but some species of microalgae are capable, if conditions permit, of producing four, eight, sixteen, or even more daughter cells in a single cell cycle^27–29^. This process, known as multiple fission^30^, is particularly studied in three green microalgae ^11^: *Parachlorella* ^31,32^ *Desmodesmus* ^33,34^ and *Chlamydomonas* ^35–37^. Their rapid growth, ease of cultivation, and ease of synchronization make them ideal model organisms for studying the eukaryotic cell cycle. These types of algae grow autotrophically during the day and can undergo reproductive sequences (DNA replication, nuclear and cellular division) at night, making the alternation of light and dark periods an effective method for cell synchronization, a crucial tool for studying their cell cycles ^30^.

Synchronization has been achieved since 1953 in *Chlorella* ^38^, and since 1972 in both *Scenedesmus* ^39^ and *Chlamydomonas* ^40^. The CC in *Chlamydomonas* consists of a variable-duration G1 phase, during which cell growth occurs and CP is attained, followed by n rounds of S/M phases to produce 2^n^ daughter cells, with each round lasting 30–40 minutes ^41^.

During the eukaryotic G1 phase, there is a conserved life cycle point-of-no-return known as START in yeast, the restriction (R) point in mammals, or the commitment point (CP) in microalgae, after which the cell will divide even in the absence of energy or nutrients. In green algae, cell size appears to be the primary prerequisite for CP attainment, which occurs when the cell reaches a critical size. Although light affects the growth rate and thus the rate at which critical cell size is reached, the critical cell size remains the same across a wide range of light conditions ^45^. CP attainment in *Chlamydomonas* is governed by two distinct but interacting pathways ^120^: 1) the retinoblastoma (RB) tumor suppressor pathway ^25^, with the RB homolog Mat3 suppressing CP attainment and E2F/DP activation promoting the expression of cell division-associated genes ^120^; and 2) CDKA1/CDK1, which is required for promotion into the division-competent state ^120^.

The algae cell cycle can be divided into a pre-CP period, which consists of the initial growth phase, and a post-CP period, which includes the overlap of one to several growth and reproductive phases (i.e. DNA replication and cell division). The CC is highly dependent on light and temperature ^37^, as first described in synchronized cultures of *Chlorella ellipsoidea* by Morimura in 1959 ^42^. This study was followed by research on other algae species ^35,43–48^ with similar results: a non-damaging increase in light intensity increases growth, mother cell size, and the number of released daughter cells. Higher light intensity shortens the cell cycle by accelerating the pre-CP phase, but not the post-CP phase, indicating that the first is size-based and the second is timer-based^45^.

Early transcriptomics analyses of the *Chlamydomonas* life cycle showed that approximately 50% of Chlamydomonas genes exhibit cyclic expression over the course of a day, with clusters related to ribosomes/translation, photosynthesis/light response, mitochondria, metabolism, cell cycle/mitosis, and microtubules/flagella displaying tightly timed expression during several light/dark cycles ^49^. This was confirmed by a more detailed transcriptome analysis of a highly synchronized population ^50^, which reinforced the cyclic properties of these genes and revealed specific expression patterns over time. For example, the photosynthesis cluster is highly expressed in light, while flagella/basal body genes begin to be expressed during the transition to light^50^.

This study uses physiology and transcriptomics to identify the metabolic state around the time of CP attainment. Two light conditions were used to distinguish CP-related genes from light-responsive genes.

## 2. MATERIAL AND METHODS

### 2.1. Growth conditions

*Chlamydomonas reinhardtii* strain 21 gr (CC–1690, wild type; Chlamydomonas Resource Center, St. Paul, Minnesota, USA, http://www.chlamy.org, accessed July 27, 2021) was grown in glass vessels containing mineral high salt (HS) medium 51 at 30°C, with aeration using air-CO2 (2% v/v). The vessels were illuminated by fluorescent tubes (OSRAM DULUX L55 W/950 Daylight, Milano, Italy), and the light intensity at the culture vessel surface was 500 μmol·m^−2^·s^−1^ of PAR. To synchronize population growth, alternating cycles of 12 hours light and 12 hours dark were applied for 3 days before the experiment, with cultures diluted to a constant cell number at the beginning of each light period ^52,53^. At the start of the experiment, the synchronized cell population was diluted to a starting concentration of about 1×10^6^ cells/mL and grown under continuous light (100 (LL) or 500 (OL) μmol photons m^−2^ s^−1^ (µmol m^-2^ s^-1^) of PAR) at 30°C. The cultures were bubbled with 2% (v/v) CO2 in air. Experiments were performed in biological triplicates, and cultures were sampled at regular intervals (hourly in OL and every two hours in LL).

#### 2.1.1. CP assay

Aliquots (1 mL) of the culture were placed on HS plates and incubated in the dark at 30°C until cell division was complete in the culture with the slowest division rate. The plates were then examined directly under a light microscope at 200× magnification. For each sample, the number of large single undivided mother cells and mother cells divided into 2, 4, or 8 daughter cells (at least 150 mother cells) were counted, and the percentage of each category was calculated and plotted over time.

#### 2.1.2. Cell division

Cell division was assessed in samples at the same time points as the CP assay, fixed with Lugol, and observed under a light microscope at 200× magnification. The percentages of undivided mother cells and mother cells divided into 2, 4, or 8 daughter cells were calculated and plotted against time.

#### 2.1.3. Cell volume

Cell volumes were measured using the Multisizer 4e (Beckman Coulter) in glutaraldehyde-fixed samples (final concentration 0.2%) by diluting 50 μL of fixed cells in 10 mL of 0.9% NaCl. The results were reported as the modal volume values of the cells in the sample.

#### 2.1.4. Quantum yield

The quantum yield (QY) of photosynthesis was measured as the F_V_/F_M_ ratio of chlorophyll fluorescence. Aliquots of 2 ml were taken from the culture and darkened for 30 minutes in 10 mm × 10 mm plastic cuvettes at room temperature. The cell suspension was mixed gently by inverting the cuvettes several times before measurement. Quantum yield was measured using an Aqua-Pen-C 100 (Photon Systems Instruments, Drasov, Czech Republic) set according to the manufacturer’s instructions ^54^.

#### 2.1.5. Photosynthetic pigments

Photosynthetic pigments were analyzed as described in Řezanka et al., 2016 ^51^. Ten milliliters of algae sample were harvested and centrifuged for 3 minutes at 4000 rpm. The cell pellet was resuspended in 1 mL of phosphate buffer (pH 7.7) containing a small amount of MgCO_3_ to stabilize the chlorophylls and stored at −20°C until analysis. Thawed cells were disrupted with 500 μL of zirconium beads (0.7 mm diameter) by vortexing for 10 minutes at 3200 rpm (Vortex-Genie 2, Scientific Industries Inc., New York, NY, USA). Chlorophylls were extracted with 4 mL of 100% acetone, followed by centrifugation for 3 minutes at 4000 rpm. The supernatant was transferred to calibrated test tubes, sealed with stoppers, and placed in a dark block. The extraction was repeated with 4 mL of 80% (v/v) acetone, and the supernatant volume was adjusted to 10 mL with 80% (v/v) acetone. To analyze chlorophyll content in the algal suspension, samples were measured on a UV– Vis spectrophotometer (UV-1800, Shimadzu, Kyoto, Japan) in 1 cm cuvettes at 750, 664, 647, 470, and 450 nm, using 80% acetone as a blank. Chlorophyll content was calculated from absorbance measured at 645 and 664 nm using the method from MacKinney ^55^. The following equations were used: Chlorophyll a (chl a): (12.25 × A664 nm – 2.79 × A647 nm), Chlorophyll b (chl b): (21.5 × A647 nm − 5.1 × A664 nm), and carotenoids: ((1000 × A470 nm) − (1.82 × chl a) − (85.02 × chl b)) / 198) ^55–57^.

#### 2.1.6. Dry mass

Biomass was separated from the medium by centrifuging 5 mL of the cell suspension in pre-weighed microtubes at 3000 g for 5 minutes. The sediment was dried at 105°C for at least 12 hours and weighed on an analytical balance (TE214S-0CE, Sartorius, Goettingen, Germany) ^58^.

#### 2.1.7. Nucleic acid and protein analysis

Total nucleic acids were extracted according to Wanka (1962) ^59^, as modified by Decallonne and Weyns (1976) ^60^. The DNA test was performed as described by Decallonne and Weyns (1976) ^60^, with modifications by Zachleder (1984) ^61^. The sediment remaining after nucleic acid extraction was quantified for protein content using the procedure described by Lowry and Randall (1951) 62.

#### 2.1.8. RNA sequencing

RNA was isolated using the ZiXpress Viral DNA/RNA Extraction Kit (Zinexts, cat. no. ZP02201-192) in the ZiXpress robot (Zinexts) according to the manufacturer’s instructions. Contaminating DNA was removed by in-column DNase treatment (NEB, cat. no. M0303) according to the manufacturer’s instructions. RNA fragmentation and rRNA depletion were performed using the QIAseq FastSelect kit (Qiagen, cat. no. 334315). The library was prepared using the NEBNext Ultra II Directional Library Prep Kit (NEB, cat. no. E7760) according to the manufacturer’s instructions.

All purification steps were carried out using SPRIselect Beads (Beckman Coulter, cat. no. B23319). The quality of the libraries was assessed by agarose gel electrophoresis. The samples were sent for RNA sequencing (150 PE) to Novogene.

### 2.2. Statistical analysis

All experiments were performed in three biological replicates (n = 3). Comparisons between experimental treatments were conducted using pairwise t-tests. A p-value less than 0.05 was considered significant.

### 2.3. Bioinformatics

#### 2.3.1. 2.4.1 RNA-seq analysis

Adapters were removed from the raw reads, and low-quality reads were filtered using fastp ^63^. The *Chlamydomonas reinhardtii* CC-4532 v6.1 genome and transcriptome were downloaded from Phytozome ^64^. The genome was indexed, and the cleaned reads were aligned using Salmon (v1.10.2) ^65^. A quality check was performed after alignment using fastQC (Babraham Bioinformatics). The Salmon quantification files were processed using tximport ^67^. The z-scores for each gene were calculated within each sample and represented as the median of log2(salmon count + 1) from three biological replicates. The counts were normalized using EdgeR ^68^, and differential expression analysis was performed with the DESeq2 algorithm ^69^, using a minimum log2(fold change) (FC) of 2 and a p-value < 0.05. Mapman annotations were generated using *Chlamydomonas reinhardtii* CC-4532 v6.1, and z-scores were assigned in Mapman v3.6.0RC1 ^70^.

#### 2.3.2. Pathway analysis and visualisation

Pathway analysis was performed using GSEA ^71,72^ in weighted mode with 1,000 permutations and a minimum gene set size of 5, with custom GMT files for each comparison. The GMT files were generated from GO terms extracted from the *Chlamydomonas reinhardtii* genome v6.1 on Phytozome and the functional prediction tool from the OMA webserver ^73^. Visualization was carried out in Cytoscape ^74^ using the GSEA result folders. The Cytoscape modules used were EnrichmentMap, Auto-Annotate, and Legend Creator. Other figures were created using RStudio (RStudio v2025.09.2+418; R v3.6.0+) ^75^, with in-house R scripts utilizing the Tidyverse suite ^76^, mainly ggplot2 ^77^. Heatmaps were generated using TBtools-II software (v2.423) ^78^.

## 3. RESULTS

The aim of this study was to identify the metabolic and transcriptomic states after dark to light transition and around CP in *Chlamydomonas*. To that purpose, cells were synchronized in the dark, then transitioned into two different light intensities regimes: at Optimal Light (OL) conditions (500 µmol m^-2^ s^-1^) and Low Light (LL) condition at (100 µmol m^-2^ s^-1^) and monitored every 1 or 2 hours, respectively, in constant light. This design allowed to describe the physiology of the cells over a 18 h time course and the transcriptome around CP attainment. As the metabolic rate was faster in OL compared to LL, different timepoints, corresponding to biologically equal timepoints in CP attainment, were used for the transcriptomic analyses. For clarity, we define a mother cell as a cell that started the experiment at 0 h, and a daughter as a cell that arose from the mother cell and define the start of the cell cycle as the time when the cells were put to light at 0h.

### 3.1 Effects of light intensity on growth kinetics

#### 3.1.1 Cell growth and division

For ease of representation, timepoints are shown for every two hours for both conditions. Half of the cells grown on OL reached CP within 1 h against 8 h in LL (Fig 1A). Cells in OL increased their cell size about 6-fold (Fig. 1D) and mostly divided to 8 daughter cells. The cells in LL grew about 3.5-fold (Fig 1D) with division into 4 daughter cells. Half the cells divided around the 15 h mark post illumination in both conditions (Fig 1A). While it has been accepted that the timer was temperature dependent ^30^, a recent study showed that the mitotic sizer triggering the S/M phase is independent of the growth condition including nutrient supply, temperature, light intensity and light duration, and thus must be different from the CP sizer and is yet to be determined ^79^.

**Figure 1:**
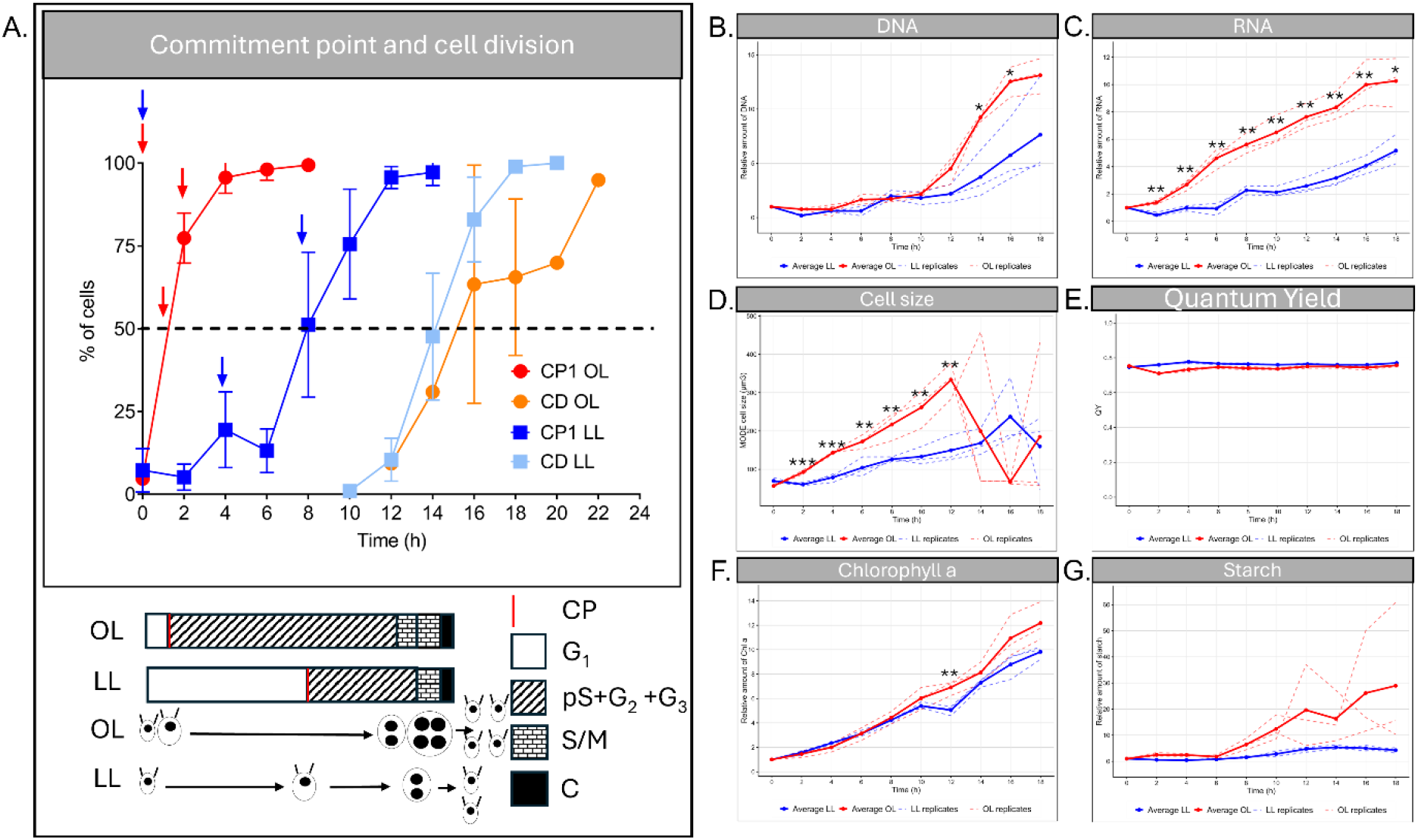
Physiology data of *Chlamydomonas* cells grown in OL and LL. **A.** Cell cycle graph of synchronised cells growing in OL and LL. The curves represent the percentage of cells attaining CP (red for OL, dark blue for LL) and dividing (orange for OL, light blue for LL). Arrows represents the sampling points for the RNA sequencing analysis. Data are represented by mean +/- standard deviation. Below is a schematic of the cell cycle in OL and LL conditions scaled on the experiments timepoints. CP = Commitment point; G1 = growth phase; pS = pre-replication phase; G2 = growth phase; G3 = growth phase between nuclear and cellular division; S/M=DNA replication and mitosis; C=cell cleavage. **B-G.** Physiological graphs of relative amount of DNA, relative amount of RNA, cell size, Quantum Yield, relative amount of Chlorophyll a, and relative amount of starch in OL (red) and LL (blue). Dotted lines are biological replicates, and bold lines represent their averages. The relative amounts were calculated against the first timepoint of the respective conditions. Asterisks represent a statistical difference at the same timepoint between the two conditions (pairwise t-test, pvalues < 0.05).

#### 3.1.1. Cell content

As the cells were smaller in LL, their content was also overall reduced. The level of RNA (Fig. 1C, S1B) and protein (Fig. S1C) was higher in OL during the whole experiment. However, the level of DNA per cell was similar between the two conditions for the first 10 h until DNA replication started (Fig. 1A). Twelve hours post-illumination, the DNA level rose in both conditions, but faster and in larger quantity in OL with a relative amount of 13.12 compared to 7.6 in LL at the maximum timepoint measured, correlating with the number of daughter cells produced in each condition (Fig. 1A).

Overall, the QY was very similar between OL and LL at all measured timepoints, indicating that the PSII was not stressed by any of the two light intensities (Fig. 1E). The levels of chlorophyll a, chlorophyll b and carotenoids are slightly (but not significantly) higher in OL compared to LL, which indicates that the LL intensity was not stressful enough to trigger changes in photosynthesis machinery at 30°C (Fig. 1F, S1F, S1G, S1H). In OL, starch was accumulated from the 8 h timepoint increasing until CD when it was rapidly consumed, shown by the huge variability intra-condition, while there was almost no starch accumulation in LL (Fig. 1G, S1H). Starch production is dependent of photosynthesis and thus light intensity, with higher (non-damaging) light intensity increasing the starch production ^80,81^.

### 3.2. RNA seq

For RNA sequencing (RNAseq), samples were taken, in 3 biological replicates, at 3 different timepoints corresponding to the dark to light transition (D/L), pre-CP1 and post-CP1. The timepoints in OL were at 0 h, 1 h, and 2 h and in LL at 0 h, 4 h, and 8 h for D/L, pre-CP1 and post-CP1 respectively (Fig. 1A). Because CP was reached rapidly in OL, we later considered the D/L sample at 0 h as D/L/pre-CP1, the sample at 1h, post-CP1 and the sample at 2h, an advanced post-CP1 (Adv_post-CP1).

In the PCA, the samples were clustering together according to their specific conditions, with only one outlier in LL post-CP1 corresponding to a sample with half the average read counts (Fig. 2A). Side analysis comparing the DESeq2 results with and without that sample showed no significant differences (>94% correlation on the log_2_(FC)) in the DEGs and the sample was kept for statistical power (Fig. S2). The samples were going in a temporal direction of D/L/pre-CP1, post-CP1, Adv_post-CP1 from top to bottom on the PC2 in OL and in D/L, pre-CP1 and post-CP1 from left to right on the PC1 in LL. The post-CP1 and Adv_post-CP1 in OL were close to each other showing that the gene expression seems to be stable after reaching CP and different to D/L/pre-CP1. In contrasts, in LL, the sampling points at 0 h, 4 h and 8 h allowed the cells to develop a more diverse transcriptome and allowed a better separation over the PC1 axis.

**Figure 2:**
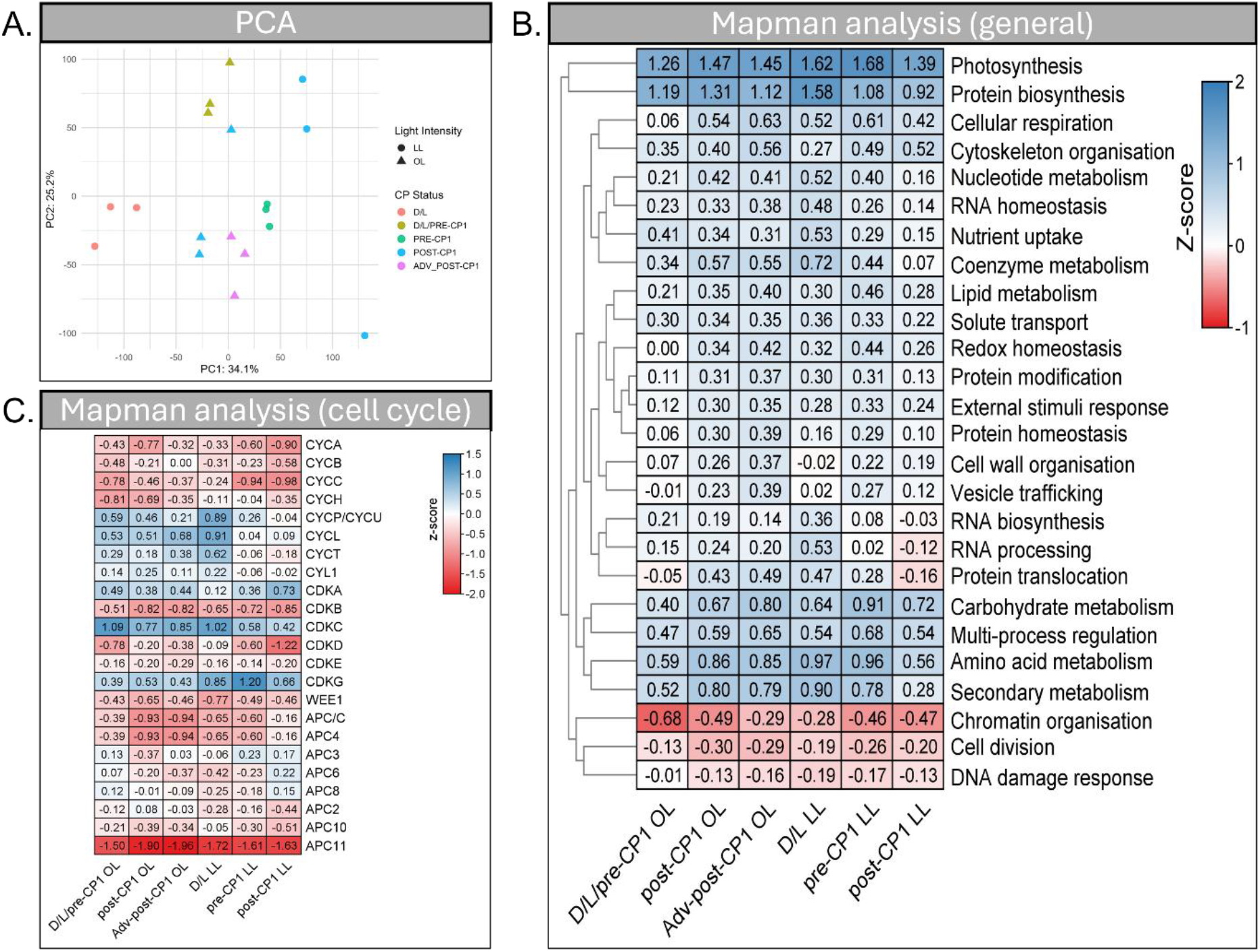
RNA-seq overview. **A.** Principal Component Analysis (PCA) of the RNA-seq samples. The first principal component (PC1) accounted for 34.1% and the second principal component (PC2) for 25.2% of the total variance of the dataset. The shapes differentiate the different light intensity, with triangles for OL and cercles for LL. The colours differentiate the biological timepoints. **B.** General Mapman analysis of the transcriptomics. Genes were assigned by Mapman to categories and their expression represented as z-scores. Red represents lowly expressed genes and blue more expressed genes. **C.** Mapman analysis of the known genes involved in the cell cycle. Red represents lowly expressed genes and blue more expressed genes.

Preliminary Mapman analysis of the transcriptomics revealed that “photosynthesis” and “protein biosynthesis” were the most expressed clusters, while “chromatin organisation” and “cell division” were the least expressed in our dataset (Fig. 2B). The “photosynthesis” cluster was the more expressed at the beginning of the cell cycle in both OL and LL with a z-score of 1.26 and 1.62 respectively. In LL, the z-score for that cluster was always higher than in OL at all measured timepoints, indicative of the need to produce more genes related to that cluster with lower light intensity, even though it was not correlated with the production of chlorophyll which was similar in OL and LL at the sequenced timepoints. In OL, this cluster was more expressed in post-CP1 compared to pre-CP1 while it was the opposite in LL, which may indicate that the gene expression related to photosynthesis may not be a criterion for CP attainment or that the minimal expression requirements for photosynthesis was lower than in our dataset. Similarly, the “protein biosynthesis” cluster was more expressed in LL at the beginning of the cell cycle compared to OL and the second most expressed cluster in the two conditions. Inside this cluster, the “ribosome biogenesis” was the most expressed sub-cluster which may hint at the importance of the ribosome in the cell cycle.

On the other hand, the “chromatin organisation” was low expressed overall in our dataset but appeared to increase over time in OL. This cluster was expected to be more expressed towards the end of the cycle, in S phase, when the DNA is duplicated and compacted before the division needing more histones than at the beginning of the cycle ^82–84^. It was shown that the production of histones starts at the beginning of the human cell cycle, with a peak of expression before the S phase concomitantly to the DNA replication possibly explaining why we observed low, but not null, expression of the “chromatin organisation” in our dataset ^85^. The low expression of the “cell division” cluster was expected because our sampling points were preceding cell division by at least 5 h. The cluster contains genes involved in the “cell cycle organisation” including CDKs and APCs, known to be an active part of the cell cycle regulation ^24^ (Fig.2C). All APCs had either very low or negative Z-scores with APC11 being the lowest with a Z-score between -1.5 and -1.96. Their low expression at the beginning of the cell cycle was previously shown and the APCs only starts to be expressed before mitosis to trigger the mitotic exit and downregulate CDKs in the following G1 phase in the daughter cell ^50^. APC/C, APC3 and APC6 have their Z-scores increasing to positive level in post-CP1 in LL which could correspond to the start of the next step of the cell cycle as this timepoint was the closest to S/M phase of the entire dataset. This increase was not quite clear in Adv_post-CP1 because the sampling was done in a still early post-CP1 (4h in OL compared to 8h in LL, closer to cell division). Most CDKs had negative or very low Z-score (*CDKB*, *CDKD*, *CDKE*) in our dataset, as previously described ^86^. Three CDKs, *CDKA1*, *CDKC1*, and *CDKG1*, had a positive Z-score. The expression of *CDKA1* in OL appeared unchanged over time due to the short temporal distance between the sampling timepoints, its level in LL increased over time with low expression in D/L (0.12) and higher in post-CP1 (0.71). This pattern was already described in Bisova et al., (2005) with *Chlamydomonas* grown at 250 µmol m^-2^ s^-1^. *CDKC1* was the CDK with the highest Z-score at 0 h in both OL and LL. In the same way as *CDKA1*, its level remained stable over sampling times in OL. However, in LL, its expression decreased by roughly half over the 8 h time course. The only publication experimentally testing its expression showed that *CDKC1* was constitutively expressed during the cell cycle ^86^.

Differential gene expression analysis was performed, comparing the gene expression between biological timepoints intra- and inter-conditions. The different comparisons are reported in Table 1. The upset plot represents the number of shared and unique genes after comparing the same biological timepoint between OL and LL (Fig 3B). Because the first sampling point in OL represents D/L/pre-CP1, the comparison was made twice, once against D/L in LL and once against pre-CP1 in LL. Thus, the comparison between D/L/pre-CP1 in OL and D/L in LL was described as [D/L]/pre-CP1 for OL, and the comparison between D/L/pre-CP1 in OL and pre-CP1 in LL was described as D/L/[pre-CP1] for OL.

**Figure 3:**
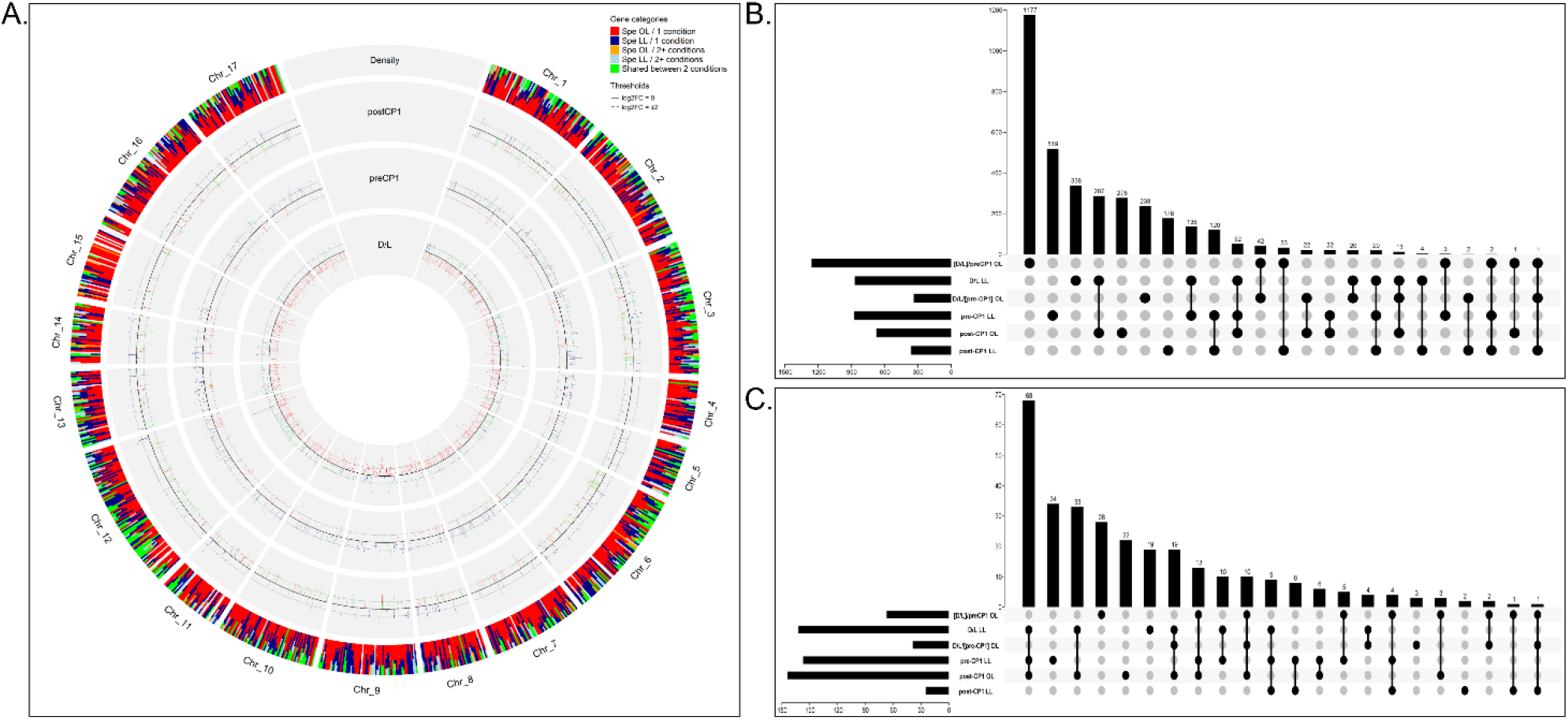
Overview of the differentially expressed genes in OL and LL. **A.** Circos plot ^87^ of the gene expression under OL and LL. From the inner track to the outer track: DEGs in OL in negative against LL in positive with their relative gene expression, in order D/L, pre-CP1 and post-CP1; density of genes on the chromosome by slice of 10000 bp. The colours represent the specificity and commonalty to OL and/or LL according to their biological timepoints. Red = specific to OL in one biological timepoint dark blue = specific to LL in one biological timepoint; orange = specific to OL and expressed in at least 2 biological timepoints; light blue = specific to LL and expressed in at least 2 biological timepoints; green = shared between OL and LL at any biological timepoints. **B.** Upset plot of the differentially expressed genes. The comparisons were made by comparing the same biological timepoint between LL and OL. Because the first sampling point in OL represents D/L/pre-CP1, the comparison was made twice, once against D/L in LL and once against pre-CP1 in LL. Thus, the comparison between D/L/pre-CP1 in OL and D/L in LL was described as [D/L]/pre-CP1 for OL, and the comparison between D/L/pre-CP1 in OL and pre-CP1 in LL was described as D/L/[pre-CP1] for OL. **C.** Upset plot of the expressed GO terms. The comparisons were made by comparing the same biological timepoint between LL and OL. Because the first sampling point in OL represents D/L/pre-CP1, the comparison was made twice, once against D/L in LL and once against pre-CP1 in LL. Thus, the comparison between D/L/pre-CP1 in OL and D/L in LL was described as [D/L]/pre-CP1 for OL, and the comparison between D/L/pre-CP1 in OL and pre-CP1 in LL was described as D/L/[pre-CP1] for OL.

**Table 1:**
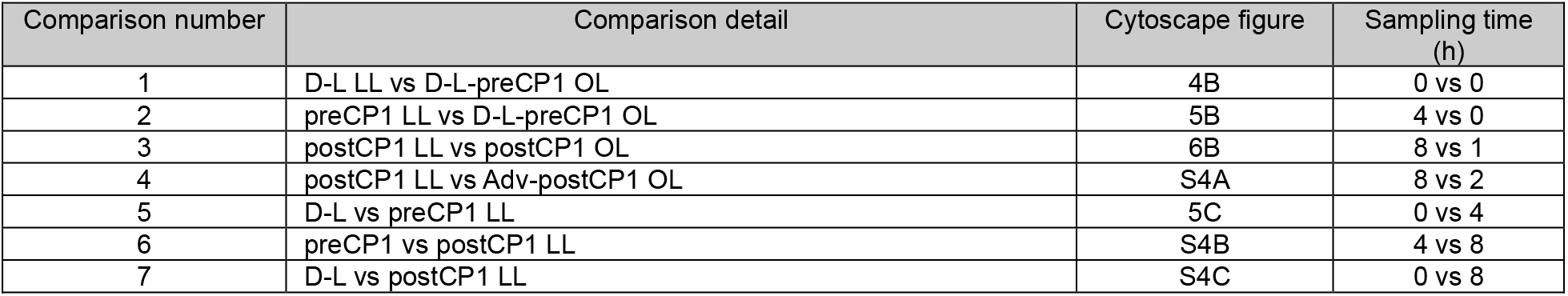
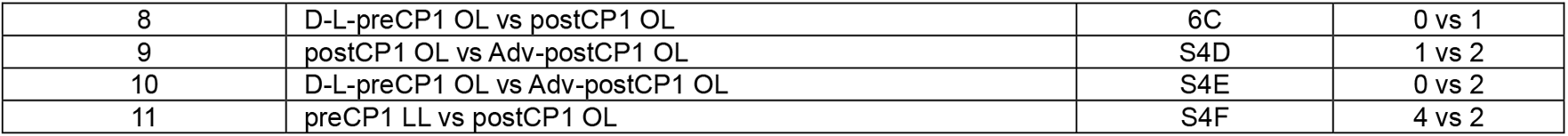
Details of the different comparisons used for the different analysis, their respective Cytoscape figure number and sampling time in hour.

The highest change in differential expression was happening during the transition from dark to light between OL and LL with 1259 genes in [D/L]/pre-CP1 in OL and 870 in D/L in LL, with 1177 genes specific to [D/L]/pre-CP1 in OL and 338 genes specific to D/L in LL. The differential expression remained high in pre-CP1 in LL when compared to D/L/[pre-CP1] with 874 genes differentially expressed in LL with 519 specifics. In OL, the second peak of differential expression was at post-CP1 compared to its LL equivalent with 675 genes differentially expressed in OL compared to 360 in LL. 278 genes were specific to post-CP1 in OL and 178 in post-CP1 in LL. Overall, in both conditions, the highest level of differential expression was at 0 h right after the transition from dark to light, with more genes differentially expressed in OL compared to LL (Fig. 3A-B). This could be triggered by one or several light switch signal(s) expressed in the dark (or constitutively expressed) ^88,89^.

In terms of gene ontology (GO), using the same comparisons and having at least 5 genes in a GO term to be acknowledged, the highest number of shared GO terms was between post-CP1 in OL and D/L and pre-CP1 in LL (Fig. 3C). The GO terms mainly composed of photosynthesis, amino acid synthesis, mitochondria, protein modifications and protein transport suggesting that those were independent of CP attainment and were probably time related as the cells were still relatively early in their development (≤4h). Overall, the number of GO terms contrasts with the number of genes expressed, probably because of the still poor gene annotation in *Chlamydomonas* where most genes don’t have a GO term or did not pass our 5 genes per GO term threshold (for example less than 246 genes out of 1177 of the specific in D/L/pre-CP1 OL compared to D/L LL were assigned to a GO term with the minimum threshold). The details of those genes and their respective GO terms are detailed in the following sections.

### 3.3 Gene Ontology analysis

To have a broader idea of the effects of light intensity on the cell cycle, multiple GSEA analysis and clustering via Cytoscape were performed. A different view of the difference of expression can be observed when comparing directly the same cell cycle phase between OL and LL. It must be noted that having genes or GO terms enriched in OL does not mean that they are not in LL, and inversely. Those could always be constitutively expressed for metabolism purposes but could be even more expressed in one of the two conditions. For example, the flagellar gene FAP380 is more expressed in D/L in LL compared to D/L/pre-CP1 in OL with a log_2_(FC) = 2.03, but it is highly expressed in both conditions with a z-score of 1.48 in OL and 1.99 in LL at 0h. The details of the Z-scores and DEGs are in supplemental table S1. The details of the GO terms are in supplemental table S2.

#### 3.3.1.1 Let there be light

Transition from dark to light is a major shift that allows the future mother cells to start to grow ^cro,90^. To identify the shared genes between D/L samples in OL and LL, we compared the gene expression and assigned GO terms in 2 comparisons: 1) D/L/pre-CP1 against post-CP1 in OL (comparison 8) and 2) D/L against pre-CP1 in LL (comparison 5). The comparison D/L/pre-CP1 in OL against D/L in LL (comparison 1) was used to determine the specific genes expressed and GO terms enriched. 126 genes were shared in D/L with a *gProfiler* ^91^ enrichment in nucleic acid binding, ribosome biogenesis, RNA processing and gene expression.

Those genes were distributed in 29 pathways shared at the 0 h timepoint in OL and LL (Fig. 4A). Among them, the terms “nucleolus”, “rRNA processing” were the most enriched ranking first and second in both OL and LL, followed up by “ribosome biogenesis”. 21 genes were shared in the GO term “nucleolus” with many being computationally predicted to be involved in the ribosome biogenesis such as Cre16.g679450 (small nucleolar RNA-associated protein 24), or Cre16.g683793 (similar to NOP7 in *S. cerevisiae* ^92^). Other enriched genes enriched at D/L in OL and LL included RNA helicases (Cre02.g118300, Cre12.g513700, Cre12.g526850) and RNA binding PUF genes (PUF1 and PUF4). PUF proteins have been more extensively studied in Opisthokonts than plants and barely in algae, but they have important roles in cell division, differentiation and development by controlling gene expression post-transcriptionally ^93–95^. PUF1 and PUF4 were found to be upregulated in *Chlamydomonas* cells grown mixotrophically in spatialised structured 3D environment, but no other studies are available to our knowledge ^96^. Overall, after the cells were transitioned from dark to light, they appeared to primary focus on protein production by transcribing genes related to ribosome biogenesis and the transcription. Comparatively, the transition from dark to OL triggered a less diverse response than in LL despite having 29% more genes expressed. This could be either consequence of the culture history, the cells were acclimated in OL before dark, or due to lower energy provided by lower light intensity in LL, which might force the cells to trigger a more complete response. Alternatively, the genes did not pass our GO terms threshold.

**Figure 4:**
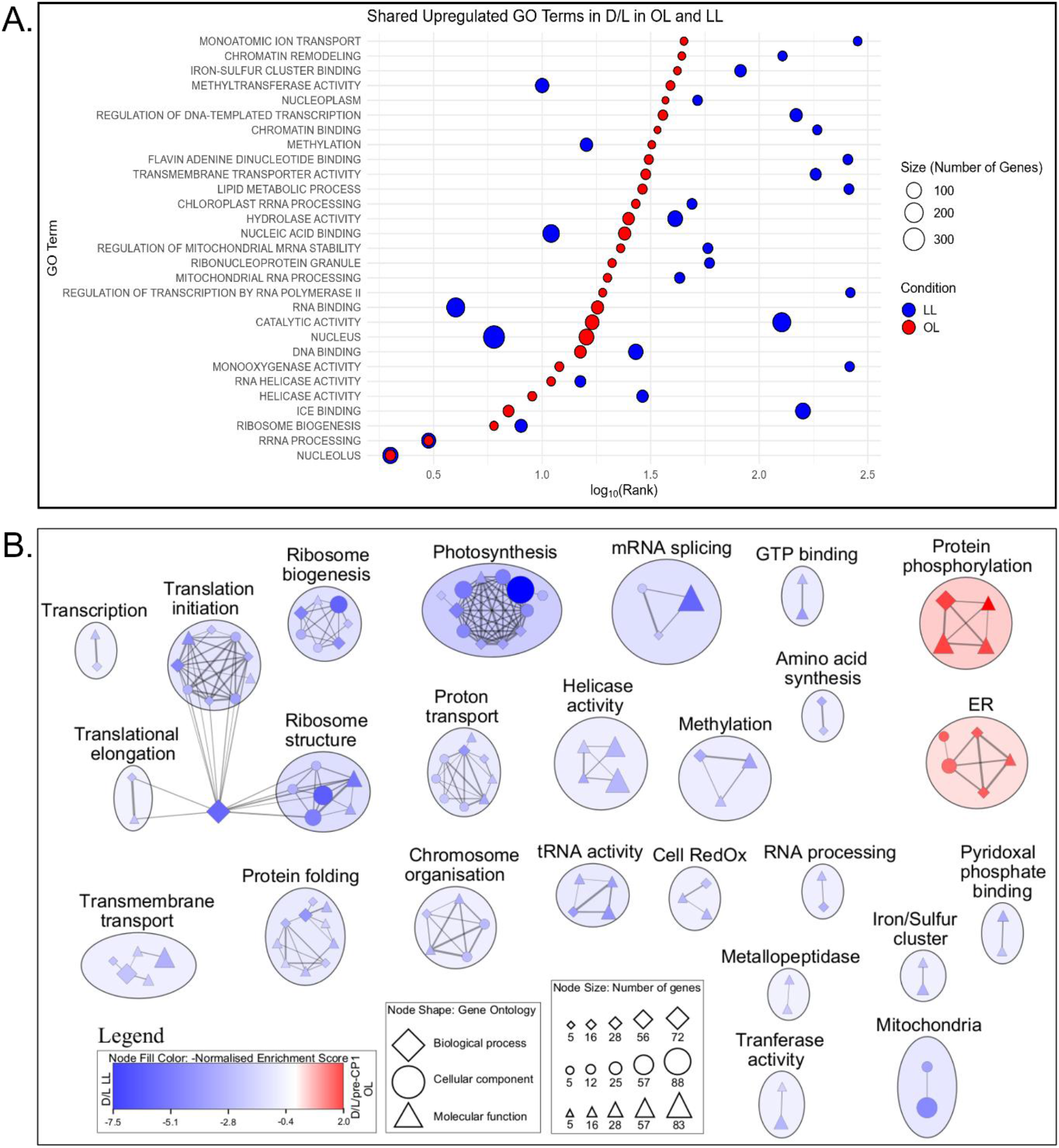
GO terms expression in D/L/pre-CP1 in OL and D/L in LL. **A.** Bubble plot of the shared GO terms upregulated in D/L in OL and in LL. Red represents OL samples and blue LL samples. The cercle size represents the number of genes in the GO term category. The rank shows the most important GO terms from left to right and was log2 transformed to fit. **B.** Network landscape of the GO terms differently enriched between D/L/pre-CP1 in OL and D/L in LL. The figure was made in Cytoscape from GSEA results using a p-value<0.01, q-value<0.05 and a similarity cutoff Jaccard+overlap coefficient of 0.375. Nodes represent GO terms and clustered together. The single nodes and the nodes’ names have been removed for ease of reading and the details are in supplementary table 2. The red color represent D/L/pre-CP1 OL, blue represents D/L LL. The symbols represent the GO categories with diamond as biological process; cercle as cellular component and triangle as molecular function. The size of the symbols represents the number of genes in the node.

Specifically, in D/L/pre-CP1 in OL compared to D/L in LL, the 5 most enriched GO terms, out of 67, are related to cytoskeleton or kinases (Fig. 4B). The “cytoskeleton” GO term was mainly composed of flagella related genes such as RSP14, FAP198, POC3 or VLF1^97–100^. Among the 60 kinases differently expressed in OL compared to LL, 3 predicted MAP kinases belong in the 5 most differentially expressed kinases: Cre11.g801238, Cre11.g467589 and Cre11.g801232 with log_2_(FC)=-6.44; -5.6 and -4.58 respectively. MAP kinases in *Chlamydomonas* play significant roles in lipid synthesis, stress response and flagellar assembly among other roles ^101–103^. These 3 genes do not have a proven function yet, alongside the other 7 MAP kinases found in this comparison (Cre03.g158300, Cre11.g801236, Cre17.g713750, Cre11.g467584, Cre02.g087850, Cre11.g801233 and Cre17.g710600), but considering the enrichment of the cytoskeleton in OL, we can hypothesize that they actively participate in the flagella assembly. 5 cyclin-dependent kinases-like (CDKL) were more expressed in D/L in OL compared to LL: Cre02.g095300, Cre01.g052850, Cre03.g187300, Cre02.g095099, Cre10.g432150. CDKL are known for participating in cell cycle regulation, but also in flagellar length and assembly ^104,105^. As they were not upregulated in D/L samples in LL, we can hypothesize that they participate in the regulation of the flagella formation, as soon as they are put in OL and later in the cell cycle in LL. 45 other kinases were differentially expressed in OL compared to LL at 0h, mainly unnamed Serine/Threonine kinases. This class of kinases have very diverse regulatory roles in the cell, such as light response, sulphur regulation, flagella formation ^106–108^. The *gProfiler* enrichment of the 1177 specific genes expressed in D/L/pre-CP1 in OL compared to D/L in LL showed an enrichment in DNA metabolism (recombination, helicase activity, integration, damage response, repair and catabolism), lipid catabolism but sphingolipids and ceramide biosynthesis involved in the membrane structure and the COP9 signalosome. The COP9 signalosome is a conserved protein complex that regulates a family of cullin-RING E3 ubiquitin ligase complexes (CRLs), and plays diverse roles in the gene expression, cell proliferation, cell cycle in animals and plants, but also the plant development in response to internal and external signals like phytohormones, light or temperature ^109–112^. While the role of the CRLs in lipid biosynthesis has been shown in *Chlamydomonas* ^113^, nothing was published on the COP9 signalosome, which may play a role in the cell cycle under optimal conditions.

At 0h, in LL compared to OL, the most expressed GO terms focused on nuclear and chloroplastic ribosomes biogenesis with emphasis on translation and photosynthesis. While cells in both conditions were expressing the same genes related to ribosome biogenesis and translation, it was more pronounced in LL. To compensate the reduced amount of photoenergy received, the cells needed to increase their protein output by increasing the level of ribosomes and translation machinery. The GO term “translation” was mainly constituted of translation initiation factors like EIF2A/G or EIF3A-K ^114^ and mitochondrial and chloroplast subunits like MRPL10/23/28 ^115^ and PRPL1-29 ^116^. Surprisingly, genes related to photosynthesis are more expressed in LL compared to OL, although the QY and the amount of chlorophyll was similar in both conditions at all measured timepoints. While the z-scores related to photosynthesis showed that this cluster was always the most expressed in our dataset, the photosynthetic genes were not as highly expressed in OL compared to LL until post-CP1 and not even reaching the z-score in D/L in LL. This indicate that the photosynthetic apparatus may not be involved in attaining CP1. However, two kinases were more expressed in LL compared to OL, Cre07.g800863 and *STL1* (Cre12.g483650) with log_2_(FC)=

2.83 and 2.93 respectively. *STL1* is a thylakoid protein kinase responsible for the regulation of photosynthesis, especially the light harvesting complex II (LHCII) ^117,118^. On the other hand, Cre07.g800863 has not been described and is unique to *Chlamydomonas*. The only information is its predicted localization to chloroplast by Predalgo ^119^. Its expression was increasing over time while comparing to OL equivalent biological timepoint with log_2_(FC)=4.31 in pre-CP1 and 4.57 in post-CP1. Taken together and with the emphasis of photosynthesis in LL compared to OL, we can assume that Cre07.g800863 is also involved in the regulation of the photosynthesis in LL by phosphorylating yet unknown targets, and further analysis should be performed to determine its precise role. 338 genes were specific to D/L in LL in the upset plot (Fig. 3B) with a *gProfiler* enrichment in the central dogma of biology with RNA polymerase and translation initiation, but also methylation (macromolecules, RNA, rRNA) and rRNA processing were enriched GO terms. The chromosome structure and the nucleosome assembly were also found to be specific to D/L in LL compared to all other comparisons (Fig. 3C). Overall, the transcriptome indicated that the metabolism was slower in LL than OL, with more central biology and photosynthesis related genes expressed.

#### 3.3.1.2 Pre-Commitment transcriptomes

During their growth in G1 phase, the cells reach CP, triggered by an unknown mechanism, probably via a sizer ^120^. Our sampling design assumed genes specific expressed at CP1 will be related to this mechanism and possibly interact with the CDKA and RB/E2F/DP CP pathways responsible for CP attainment ^121,122^. To determine the genes differently expressed in pre-CP1 compared to the rest of the cell cycle, comparison of D/L/pre-CP1 against post-CP1 was used for OL conditions, and two comparisons were combined and used for LL conditions: D/L against pre-CP1 (5) and pre-CP1 against post-CP1 (6).

21 GO terms were shared at pre-CP1 between OL and LL with more variability in the GSEA ranks compared to D/L (Fig. 5A). Among the shared, in OL, the focus was on protein phosphorylation. Most pathways ranked higher in LL, so less important in all the GO terms from their respective comparisons. This could reflect the time difference in sampling and the faster metabolism in OL, where more genes were expressed, while in LL, gene expression might be more specific.

**Figure 5:**
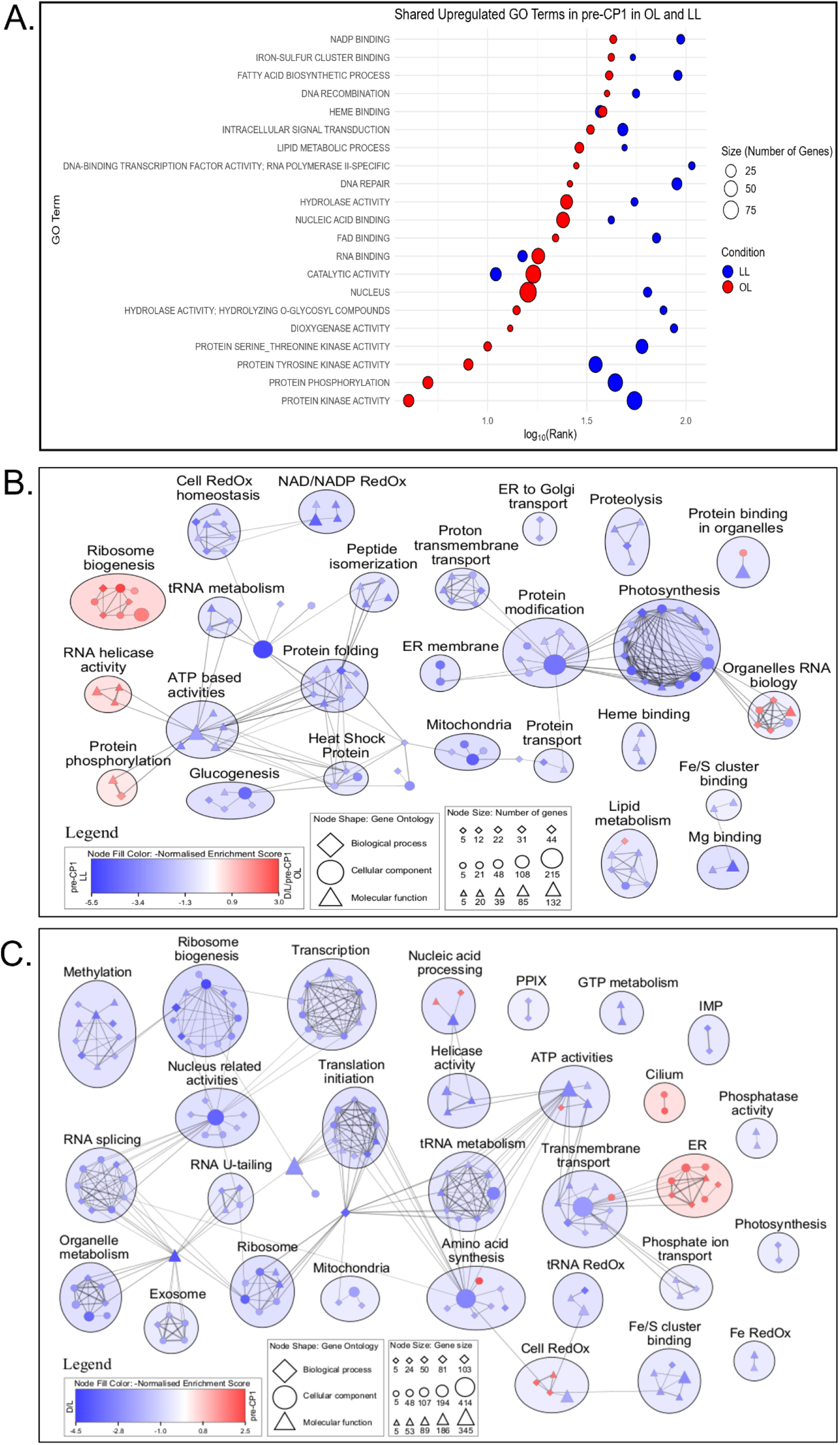
GO terms expression in D/L/pre-CP1 in OL and D/L in LL. **A.** Bubble plot of the shared GO terms upregulated in pre-CP1 in OL and in LL. Red represents OL samples and blue LL samples. The cercle size represents the number of genes in the GO term category. The rank shows the most important GO terms from left to right and was log2 transformed to fit. **B.** Network landscape of the GO terms differently enriched between D/L/pre-CP1 in OL and pre-CP1 in LL. The figure was made in Cytoscape from GSEA results using a p-value<0.01, q-value<0.05 and a similarity cutoff Jaccard+overlap coefficient of 0.375. Nodes represent GO terms and clustered together. The single nodes and the nodes’ names have been removed for ease of reading and the details are in supplementary table 2. The red color represent D/L/pre-CP1 OL, blue represents pre-CP1 LL. The symbols represent the GO categories with diamond as biological process; cercle as cellular component and triangle as molecular function. The size of the symbols represents the number of genes in the node. **C.** Network landscape of the GO terms differently enriched between D/L and pre-CP1 in LL. The figure was made in Cytoscape from GSEA results using a p-value<0.01, q-value<0.05 and a similarity cutoff Jaccard+overlap coefficient of 0.375. Nodes represent GO terms and clustered together. The single nodes and the nodes’ names have been removed for ease of reading and the details are in supplementary table 2. The red color represent pre-CP1 LL, blue represents D/L LL. The symbols represent the GO categories with diamond as biological process; cercle as cellular component and triangle as molecular function. The size of the symbols represents the number of genes in the node.

The most recently proposed model to trigger CP involves a transcription factor (TF), an inhibitor (IN) and starter kinase (SK) ^79^. During the early stage of the G1 phase, IN is inactivating TF, while during growth SK slowly inhibit IN’s activity, allowing the TF to be expressed at CP. After closer look to the kinases differently expressed in the above comparisons, only 1 out of 63 was shared between OL and LL: Cre12.g483650. However, this kinase had opposite expression depending of the light condition with more expression in post-CP1 (z-score=1.099) compared to D/L/pre-CP1 in OL (z-score=0.268), and more expression in D/L (z-score=1.299) and pre-CP1 (z-score=1.073) compared to post-CP1 (Z-score=0.074) in LL. This gene, *STT7*, is a paralog of *Arabidopsis*’ *STN7* gene, responsible for LHCII phosphorylation and required for the transition from state 1 to 2 in photosynthesis ^106,123^. Because of its role in photosynthesis and different pattern of expression, we can rule out this kinase as the SK. None of the known CDKs were differently expressed in our dataset, mainly because they were constitutively expressed and regulated by phosphorylation like *CDKA1* (z-scores: 0.49/0.38/0.44 in OL and 0.12/0.36/0.73 in LL), *CDKC1* (z-scores: 1.09/0.77/0.85 in OL and 1.02/0.58/0.42 in LL) or *CDKH1* (z-scores: 0.16/0.16/0.27 in OL and 0.53/0.23/0.06 in LL) or because they were expressed before the S/M like *CDKB1* (z-score: -0.51/-0.82/-0.82 in OL and -0.65/-0.72/-0.85 in LL) and *CDKG1* (z-scores: 0.39/0.53/0.43 in OL and 0.85/1.20/0.66 in LL) (Fig. 2C) ^86^. Similarly, no cyclins were differentially expressed in pre-CP1 in both OL and LL.

When comparing D/L/pre-CP1 in OL and pre-CP1 in LL, a shift in the clusters can be observed in OL. The ribosome biogenesis and RNA helicase activity clusters were more enriched in OL than LL. Otherwise, the expression pattern was similar in LL.

As D/L and pre-CP1 had a similar network in LL and in OL, a direct comparison between D/L and pre-CP1 in LL has been made, to determine the differently expressed genes between the light transition from dark and a more advanced growth in LL. 1730 and 1173 genes were differently expressed, distributed in 304 and 106 GO terms in D/L and pre-CP1 respectively. All the enriched GO terms discussed above in D/L/pre-CP1 OL compared to D/L LL were more enriched in D/L than pre-CP1, indicating that they were downregulated over time.

In pre-CP1, an emphasis was on the membrane structure and the cell motility with the 5 most enriched pathways being “cytoskeleton”, “axoneme”, “ceramide biosynthetic process”, “phospholipase activity”, and “cellular oxidant detoxification”. The first 4 pathways include, compared to D/L, increased of number of genes involved used for cell growth and movement, i. e. genes involved in the production of membrane lipids, microtubules and the cilium. The fifth GO term, “cellular oxidant detoxification”, was led by the 4 Hybrid Cluster Proteins (*HCP1-4*) genes with expression increasing in time (log_2_(FC) from 2.27 to 5.97 in LL in pre-CP1 and post-CP1 compared to D/L). They are metalloproteins with important RedOx capacity, characterized by the presence of an iron-sulfur-oxygen cluster and mostly located in the chloroplast, except for the mitochondrial *HCP2* ^124^. They were found to be highly expressed in nitrogen starvation, darkness and anaerobic conditions, with *HCP4* having a 1500-fold change in dark anoxia ^124–126^. Despite lacking experimental evidence, their hypothesized role is molecular switch reducing nitrate in nitrite and catalysing the S-nitrosylation of other proteins ^124^. Their role in our dataset is unclear, but considering their RedOx capacity and localisation, we propose that they can also play a role in the electron exchange in the chloroplast in lower light intensity. Genes related to the endoplasmic reticulum (ER), ranked 7 out of 106, were also more expressed in pre-CP1. On top of being the main producer and distributor of proteins in cells ^127–129^, the ER also regulates the lipid metabolism ^130^ and the stress response ^131,132^. Of ER-related genes, *CRT2* was among the most upregulated genes in pre-CP1 compared to D/L in LL with a DEG of -5.49. Its z-scores were negative at 0 h and increased in the second and third tested timepoints in both light conditions. CRTs are conserved calreticulins with a high Ca^2+^ binding capacity localised in ER of Eukaryotes ^133,134^. ER is also a major Ca^2+^ storage ^135^ and is in close contact with several organelles such as the Golgi apparatus, lipid bodies, plasma membrane and chloroplast in microalgae and plants ^136–138^. Ca^2+^ is involved in photosynthesis, flagellar movement by phototaxis, stress response and sexual reproduction ^139,140^. *CRT2* was not yet studied in *Chlamydomonas* but was proposed to be involved in gamete formation with an increased level of protein in pre-gametes and gametes ^141^. We hypothesize that it was involved in the supply of Ca^2+^ to the cell for phototaxis and photosynthesis during the growth. Among other expressed genes in D/L compared to pre-CP1 in LL, 4 members of the SEC family (*SEC22/23B/61B/61G*) were more expressed in pre-CP1 with DEGs ranging from -2.16 to -3.22. SEC (Secretory) genes are part of the ER involved in protein translocation, and protein and lipid transport to the Golgi apparatus ^129,142,143^. *SEC23* and *SEC24* are part of the COP II complex, a conserved complex involved in vesicle trafficking from the ER to the Golgi apparatus ^144^. They expression increased upon light exposure ^145^. *SEC61* complex is involved in protein translocation permitting the entry of newly formed proteins in the ER ^142,146^ and its expression follows logically the ribosome biogenesis from D/L.

#### 3.3.1.3 Post-commitment expression

As we used post-CP1 sample to determine the genes expressed before CP1 in OL and LL, we can now do the opposite to determine part of the transcriptome expressed after CP in OL and LL. Again, D/L/pre-CP1 compared to post-CP1 was used to describe the difference of gene expression in OL, while the comparison between D/L and post-CP1 was used in LL. The latter was used because only 88 DEGs were found to be upregulated when comparing pre-CP1 and post-CP1, with only 12 of them having a name, limiting the analysis. It must be kept in mind that while the cell populations are at the same biological timepoint in their cell cycle (Fig. 1A), the sampling of the cells in OL was done 1h post illumination but 8h for those in LL, which necessarily would have impacted the differential expression analysis.

125 genes were shared between OL and LL at post-CP1 with a *gProfiler* enrichment on the ER and the post-translational protein transport which were the most important shared GO terms based on GSEA analysis (Fig. 6A). Most of the other shared GO terms in Fig. 6A were ranked low in their respective comparisons. The exceptions were “cytoskeleton” and “DNA integration” ranked 1 and 4 respectively in LL but still being ranked as low as 119 and 142, respectively, in OL. This was probably an effect of time over biology as cells in OL post-CP1 were ∼14h from cell division while the cells in LL were ∼7h away and were already expressing more genes related to the cytoskeleton, like basal body and flagellar genes. The genes in DNA recombination are involved in quality control of the DNA integrity before starting the DNA replication which was in line with increasing DNA in our physiology data from ∼8h. Indeed, 10 genes out of 19 have a predicted definition (pdef) of DNA breaking-rejoining enzyme, and 2 are predicted to be DNA repair proteins.

**Figure 6:**
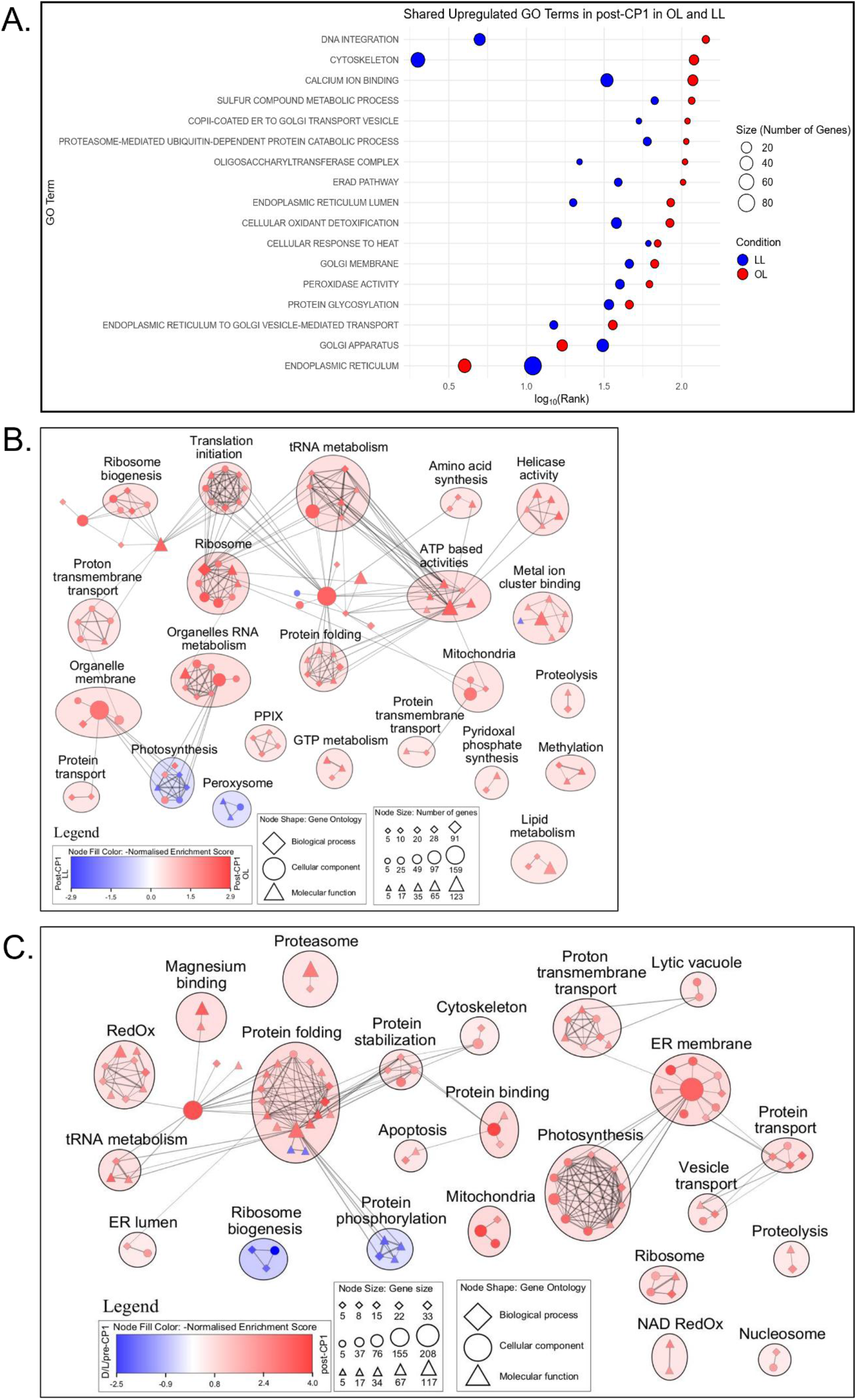
GO terms expression in post-CP1 in OL and in LL. **A.** Bubble plot of the shared GO terms upregulated in post-CP1 in OL and in LL. Red represents OL samples and blue LL samples. The cercle size represents the number of genes in the GO term category. The rank shows the most important GO terms from left to right and was log2 transformed to fit. **B.** Network landscape of the GO terms differently enriched between post-CP1 in OL and LL. The figure was made in Cytoscape from GSEA results using a p-value<0.01, q-value<0.05 and a similarity cutoff Jaccard+overlap coefficient of 0.375. Nodes represent GO terms and clustered together. The single nodes and the nodes’ names have been removed for ease of reading and the details are in supplementary table 2. The red color represent post-CP1 OL, blue represents post-CP1 LL. The symbols represent the GO categories with diamond as biological process; cercle as cellular component and triangle as molecular function. The size of the symbols represents the number of genes in the node. **C.** Network landscape of the GO terms differently enriched between D/L/pre-CP1 and post-CP1 in OL. The figure was made in Cytoscape from GSEA results using a p-value<0.01, q-value<0.05 and a similarity cutoff Jaccard+overlap coefficient of 0.375. Nodes represent GO terms and clustered together. The single nodes and the nodes’ names have been removed for ease of reading and the details are in supplementary table 2. The red color represent post-CP1 OL, blue represents D/L/pre-CP1 LL. The symbols represent the GO categories with diamond as biological process; cercle as cellular component and triangle as molecular function. The size of the symbols represents the number of genes in the node.

When comparing gene expression in post-CP1 in OL against LL, there is more expression in OL, with 675 genes differently expressed within 174 GO terms in OL against 360 genes within 25 GO terms in LL. This could reflect another time over biology effect of our sampling as CP was reached later in LL allowing the cells to develop their transcriptomes before CP in LL. Also, the number of genes expressed was decreasing over time with the dark to light transition involving a big transcriptomic adaptation (Fig. 3A-B).

After 1h in OL, the cells produced a more diverse transcriptome, still with a focus on ribosome biogenesis and translation, but also expressing genes related to photosynthesis. As the photosynthetic system is constitutively expressed in both conditions, and *Chlamydomonas* can reach CP in heterotrophic conditions ^90,147^, we can assume that may not be necessary for triggering CP, although it could indirectly accelerate it by providing faster and more abundant resources to the cell. Many other basic metabolic clusters were enriched in OL, like the organelle biogenesis (chloroplast, mitochondria), the protein processing (translation, folding, modification), the synthesis of amino acids or activating the lipid metabolism. The expression of all those genes probably reflects that the cells grew faster and divided into more daughter cells. Most of the 278 genes unique to post-CP1 in OL from the upset plot have a *gProfiler* enrichment in different steps of translation such as the amino acid metabolism, tRNA modification or the regulation of translational fidelity. The GO term “cell component” was mainly enriched in genes involved in plastids like chloroplast and mitochondria. A direct comparison between D/L/pre-CP1 and post-CP1 in OL shows that shift of expression and the activation of the metabolism with an emphasis on protein processing (folding, binding, stabilization), ER metabolism and photosynthesis (Fig. 6C).

It is worth mentioning that not much difference happened in the hour separating post-CP1 and Advanced post-CP1 in OL, with respectively 11 and 41 genes differentially expressed (Supp Fig. S3). Compared to the hour separating D/L/pre-CP1 to post-CP1 with 1198 DEGs, the transcriptome became remarkably stable only 2h into the cell cycle, even though a huge number of gene were expressed at the start of the cell cycle. Out of the 11 genes differently expressed in post-CP1, only one was named, the purine oxidase *XDH1* with a log_2_(FC)=2.47, while one outlier with log_2_(FC)=20.69 has been found. This gene, Cre11.g801266, has not yet been described but is involved in the mitochondria and specific, yet poorly conserved, to some of the *Chlamydomonadales* order. In Adv_post-CP1, mainly cellular component GO terms are enriched like membrane, cytoplasm or ER. Between the membrane and plasma membrane pathways, 3 genes are shared: The glucan synthase *GLS2* (Cre06.g302050), an LGC-12 protein coding (Cre35.g802212), and an Inositol-phosphate phosphatase (Cre08.g376100). The second gene is an ion transmembrane transporter and participates the most in those two pathways (log_2_(FC)=-9.36). Interestingly, two Light-Harvesting Chlorophyll a/b Binding protein of LHCII, *LHCBM4* and *LHCBM8*, were expressed in post-CP1 with log_2_(FC)=-2.05 and -3 respectively but highly expressed in the same way in LL with log_2_(FC)=-6.2 and -7.15. In the cytosol, gene enriched, *GMP1*, *PROB1* or *UGP1,* were related to sugar modifications, especially phosphorylation.

In LL, compared to OL, the cells continued to express genes related to photosynthesis to continuously cope with the lower light intensity. They were also emphasising on producing peroxisome to process fatty acids and develop their membranes during their growth. The *gProfiler* enrichment of the 178 unique genes showed an increase of RedOx activity, probably due to photosynthesis and a degradation of small molecules like amino acids or carboxylic acid. Apart from those, the gene expression did not change between pre-CP1 and post-CP1 in LL.

#### 3.3.1.3 Light related response

Until now, we focused on the general transcriptome of *Chlamydomonas* in different times of the cell cycle. However, we also used different light intensities to discriminate genes related to light from the core CP genes. Here, we will describe light responsive genes as genes differentially expressed at 0h, i.e. after light illumination, independently from the light intensity, with positive z-scores in both OL and LL. Then, we will define OL light sensitive genes as genes with positive z-score at 0h in OL, but negative at 0h in LL while being differentially expressed between D/L/pre-CP1 in OL and D/L in LL. Inversely, LL sensitive genes are defined as genes with negative z-score at 0h in OL, but positive at 0h in LL while being differentially expressed between D/L/pre-CP1 in OL and D/L in LL.

632 genes were assigned to light responsive genes, with a *gProfiler* enrichment mainly in translation, ribosome biogenesis, photosynthesis, and amino acid production. The ribosome related enrichment is composed of numerous nuclear translation initiation factors such as *EIF1-3* subunits and chloroplastic ribosomal subunits such as 21 *PRPL*s and 8 *PRPS*s having the highest z-scores in that category in both OL and LL. Photosynthesis related genes were composed of chloroplast precursors like PSAs, PSBs with *PSAI1* and *PSAN1* being the most expressed of the precursors with z-scores of 1.45 and 2.19 for PSAI1 and 2.16 and 2.73 for *PSAN1* in OL and LL respectively. Another highly expressed chloroplast-related gene was *THF1*, with a z-score of 2.03 in OL and 2.5 in LL, is involved in the thylakoid formation with the *Arabidopsis* mutant *thf1* showing vesicles without thylakoid membrane. The most expressed genes related to photosynthesis were also found in the GO term “response to light stimulus” and include the light harvesting proteins *LHCA/B* with z-scores between 1.48 and 3.26 in OL and 2.16 and 3.61 in LL, with all respective genes in LL having a higher z-score than in OL. The two light responsive genes with the highest overall z-scores were *tufA* in OL with a z-score of 4.06 in OL and 3.99 in LL, and *I-creI* in LL with a z-score of 3.96 in OL and 4.26 in LL. While the chloroplastic elongation factor *tufA* was previously shown to be a circadian gene, its expression is highly increased in the first moments *Chlamydomonas* is set to light conditions and decreased in the second half of the light period ^148–150^. *I-creI* is a homing endonuclease targeting DNA with a target site of 22 bp ^151–153^. No studies to our knowledge describe the targets apart from the target site, and no link has been established to light. Further study will be required to elucidate why is this gene one of the highest expressed in D/L transition and remains high in the latter tested timepoints with z-scores above 3.95.

The 440 OL sensitive genes had a *gProfiler* enrichment in microtubules and cytoskeleton, mainly flagellar genes like *BLD10*, *FAP17/181/190*, and *VFL5*. There was also an enrichment in DNA repair and damage response as well as cellular response to stress. The genes related to those enrichments had negative z-scores in LL, probably because the difference of light intensities between the dark and light phases is lower in LL than in OL. The cells in OL were not photosynthetically stressed, as shown by the QY (Fig. 1E) and the 20 genes attributed to “DNA damage response” and “DNA repair” are mainly composed of predicted DNA mismatch repair gene such as *MLH3*, and DNA polymerases like *POLQ1* and *POL1C*. Assuming that this last part appears to be bioinformatic bias, we propose that the main response to OL is to produce flagellar related and cytoskeleton proteins for growth and phototaxis.

As the cells in LL coped with lower light intensity than in OL, the 215 LL sensitive genes had a *gProfiler* enrichment focusing on RedOx activity mainly related to photosynthesis, response to photooxidative stress, and a cellular compartmentation GO enrichment around the chloroplast and its thylakoids with genes involved in the light-harvesting complex like *LHCR1*, *LHCRB1-2*, and involved in chlorophyll binding like *ELIP2/6* being more expressed than in OL. There was also an enrichment in amino acids processing, specifically in Arginine with genes like *LCI8*, *NAGS1*, *NAGK1* and *AGS1*. Arginine can act as a nitrogen (N) reserve due to a high N to Carbon (C) ratio compared to other amino acids ^154–157^ while N starved *Chlamydomonas* reduce their photosynthetic efficiency ^103^.

## 4. DISCUSSION

In the green alga *Chlamydomonas*, cell division occurs following a multiple fission model with several rounds of S/M phases happening before the division, giving birth to 2^n^ daughter cells. During growth, the cells go through CP, a point-of-no-return from where the cells can divide even if the environmental resources are removed. While the physiological basis of CP has been well described, the molecular processes that precede and potentially regulate this transition remain unclear. Our combined physiological and transcriptomic analyses provide new insights into gene expression dynamics occurring before/at CP. We initially started with a physiology approach to describe the cell cycle in OL and LL. Overall, the cells in OL were bigger and contained more of macromolecules than the cells in LL. They also give birth to eight daughter cells compared to four in LL. Despite CP was attained later in LL, the cells divided at the same time as the cells growing in OL around 15 h. That confirms that the cell division timing is not affected by cell size ^79^, but by a timer which is temperature dependent ^29,158^. As our cells were both grown at the same temperature and attained at least one CP, they were able to divide at the same time. This study mainly aimed at characterization of the gene expression around CP based on the hypothesis that genes specifically expressed before or after CP may participate in or affect CP attainment. Due to fast CP attainment in OL, within 1-2 hours after transfer from dark to light, we used two light intensities to clean the CP related dataset from light responsive genes which respond to the light on signal and would temporally correlate to CP attainment but are not functionally related.

Overall, we were unable to identify single candidate gene or set of genes that would be expressed around CP attainment and possibly related to it. This could be for several reasons: (1) The genes are not expressed just before CP1, but are rather constitutively expressed during the cell cycle, with or without a peak of expression before the S/M in the mother cell. Such protein upscaling was recently found a candidate in *Chlamydomonas*, *TNY1* localised upstream of *CDKG1*, that is passed down generation at equal level in the daughter cells. As its expression is constant, when the cell growth, it gets diluted and exert less repression on CDKG1 triggering extra divisions in the S/M phase ^159^. Other similar subscaling proteins like Whi5 and RB in budding yeast and mammals respectively, were found to modulate the regulators of the S phase ^160–162^. The canonical cell cycle regulators, cyclin-dependent kinase (CDKs) are in *Chlamydomonas* expressed constitutively and/or in higher quantity before the S/M phase ^86^. From that, we can hypothesize that they could also be targets of other upscaling proteins and the cell cycle regulators are expressed before S/M phase and passed down generation. (2) The CP attainment is multi-composite and it is not defined by genes but rather by physiological or metabolic state. Based on the focus on ribosome biogenesis in our dataset at the very beginning of their cell cycles, we propose a ribosome number/concentration model that could be tested by profiling several rRNAs (nucleolar and chloroplastic) in control conditions and in presence of ribosome biogenesis inhibitor, such as usnic acid, which was tested in yeast ^163^, rifampicin for chloroplast ^164^, or silencing the ribosome biogenesis by genetic manipulation.

Our approach allowed us to describe how the light intensity affects the transcriptome of synchronised *Chlamydomonas* cells. We observed that the LL conditions produced a more diverse transcriptome after transition from dark to light compared to OL conditions. This can be explained by: (1) Poor gene annotation and threshold of 5 genes minimum per GO terms might be too stringent and did not catch the diversity of gene expression in OL. (2) Cells in LL received less photosynthetic energy from the environment and needed to improve their translational and photosynthetic capacity to cope with that as quickly as possible. (3) Cells in LL were acclimated to OL before the dark and thus had to adapt to the new lower light intensity ^81,165^. (4) The time required to achieve CP is longer in LL, meaning that the cells had more time to develop their transcriptomes. We also show that the photosynthetic system may not be necessary to reach CP as it is constitutively expressed in both light conditions according to z-scores and *Chlamydomonas* can reach CP in heterotrophic conditions where the photosynthetic machinery is not required ^90,147^. However, photosynthesis could be indirectly involved in the cell cycle as it brings non negligeable amount of energy to the cells. Moreover, most of the gene expression is happening in the D/L samples because of the activation of the entire metabolism upon switch to light. The light intensity triggered different responses with the LL leading to a higher production of photosynthetic genes than in OL conditions. All those pathways activated by light must be under the regulation of several light responsive genes, such as *LHCRS1* and *LHCRS3*, which had the highest z-scores of the photosynthesis-related genes in both OL and LL, but more in LL ^145,166^. *tufA* could be a promising candidate as a regulator of the circadian rhythm from early light signalling due to its expression pattern ^148–150^ and its high expression could be a signal to trigger other genes. It could be also interesting to study *I-creI* from a light response perspective and define its potential targets as it was one of the most expressed genes at the start of the experiment and maintained itself at high expression.

Finally, the transcriptomic approach still has its limits in *Chlamydomonas*: there is a diversity of gene expression between the different wild type strains grown on the same conditions, which makes the comparison of different transcriptomic studies complicated ^167^. On top on that, some genes remain poorly annotated. For example, in our dataset, the GO term “ice binding” was in the top 10 of the most enriched terms in several comparisons (1, 4, 6, 8, 10). As the cells were cultivated at 30°C, even during the dark phase, annotation of the genes as antifreeze proteins does not quite fit. In the same GO term, other genes not annotated as antifreeze proteins, such as Cre06.g293100 and Cre11.g480700, are too lacking proper annotations although they were computationally inferred as “Golgi vesicle transport” and “ceramide biosynthetic process/sphingolipid catabolic process” in the previous version of the genome. In total, 657 genes were attributed to the ice binding pathway in the genome version 6.1; 235 of them were expressed in the present dataset. Those bioinformatically misannotated genes could play important physiological roles and need a better curation to understand their roles in *Chlamydomonas*.

## AKNOWLEDGMENTS

This work was supported by the GACR project no. 22-21450S and the P JAC project “Photomachines” Reg. N° CZ.02.01.01/00/22_008/0004624. Computational resources were provided by the e-INFRA CZ project (ID:90254), supported by the Ministry of Education, Youth and Sports of the Czech Republic and by the ELIXIR-CZ project (ID:90255), part of the international ELIXIR infrastructure.

## SUPPLEMENTARY MATERIALS

**Figure S1:**
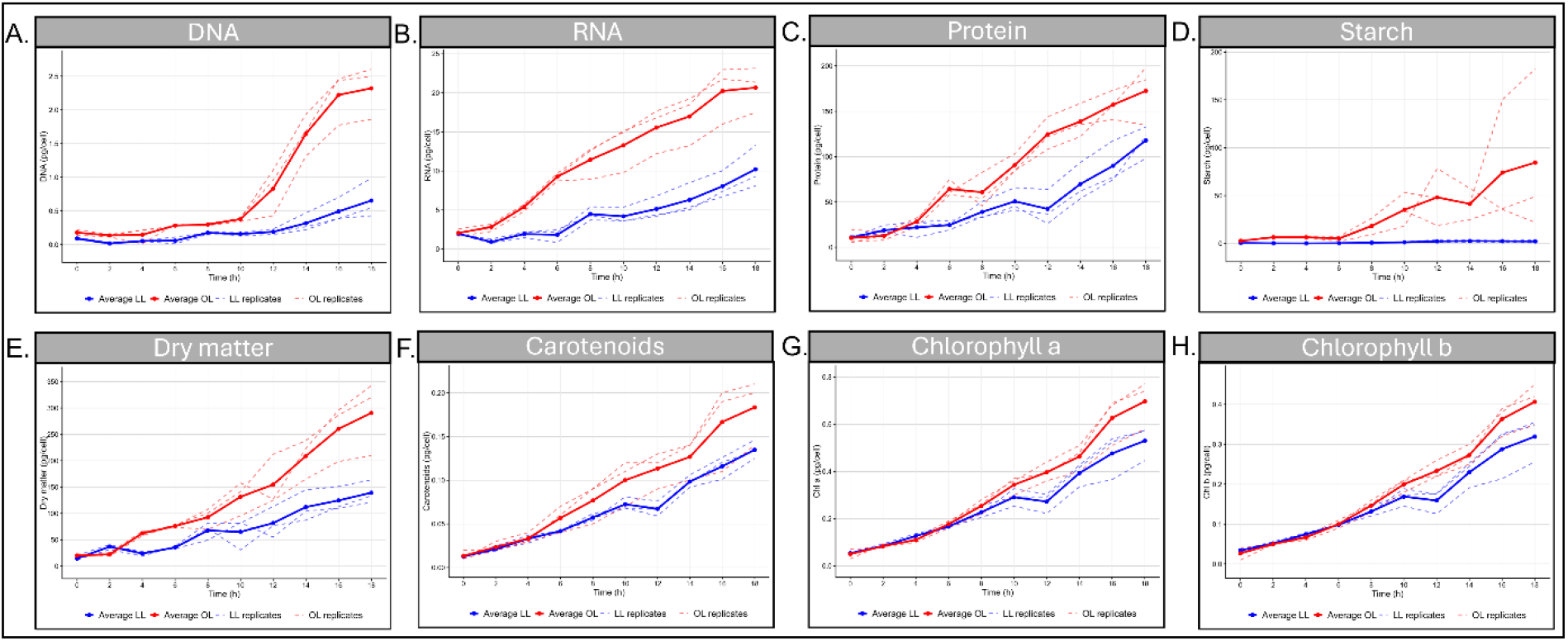
Physiology data of Chlamydomonas reinhardtii cells grown in OL and LL. **A-H.** Physiological graphs of the amount of DNA per cell, amount of RNA per cell, amount of protein per cell, amount of starch per cell, quantity of dry matter per cell, amount of carotenoids per cell, amount of chlorophyll a per cell, and amount of chlorophyll b per cell in OL (red) and LL (blue). Dotted lines are biological replicates, and bold lines represent their averages. Values are plotted in pg.cell^-1^.

**Figure S2:**
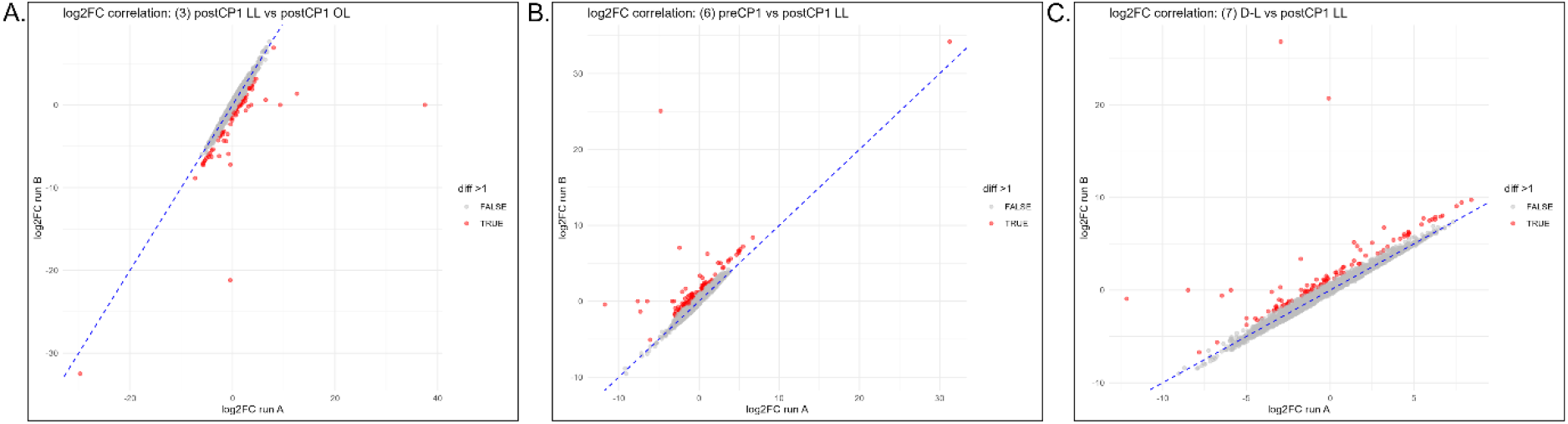
Scatter plots of DEGs excluding (Run A) or including (Run B) Chlamy17. **A.** Scatter plot of the DEGs between post-CP1 LL and post-CP1 OL. **B.** Scatter plot of the DEGs between pre-CP1 LL and post-CP1 LL. **C.** Scatter plot of the DEGs between DL LL and post-CP1 LL.

**Figure S3:**
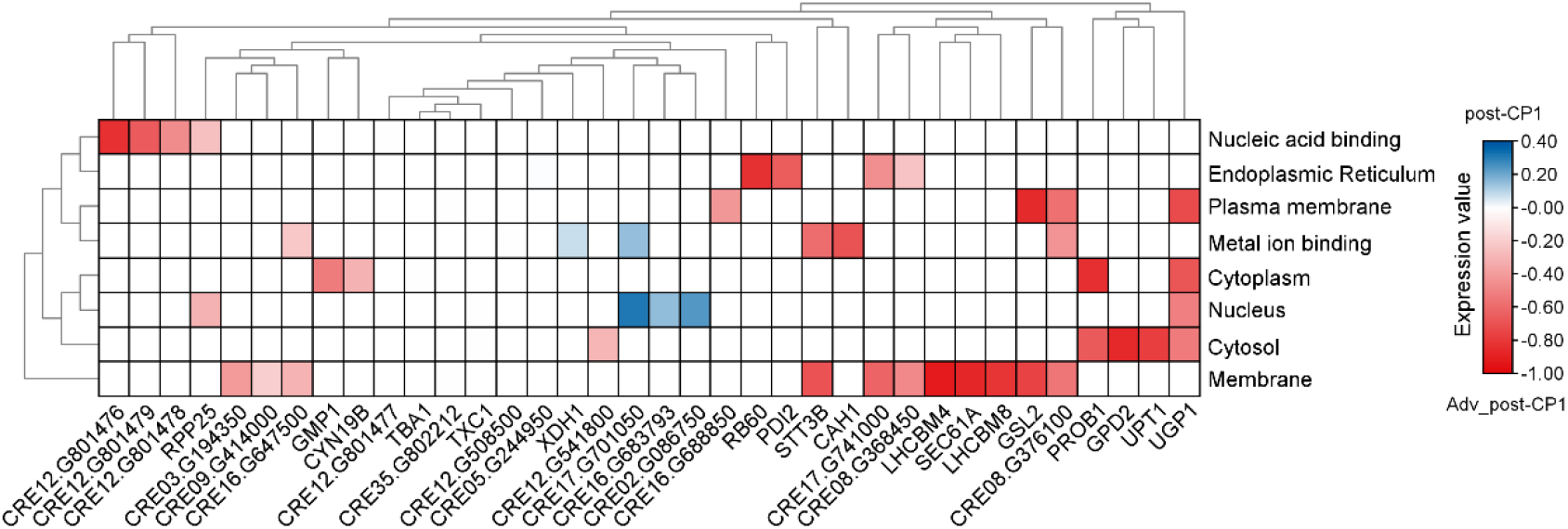
Heatmap of the DEGs between post-CP1 and Adv_post-CP1 in OL. Gene importance in a GO term is represented as an expression value calculated during the GSEA analysis. An expression value of 1 indicates the gene is in the leading-edge subset for the gene set. Red = Adv-post-CP1 and blue = post-CP1.

**Figure S4:**
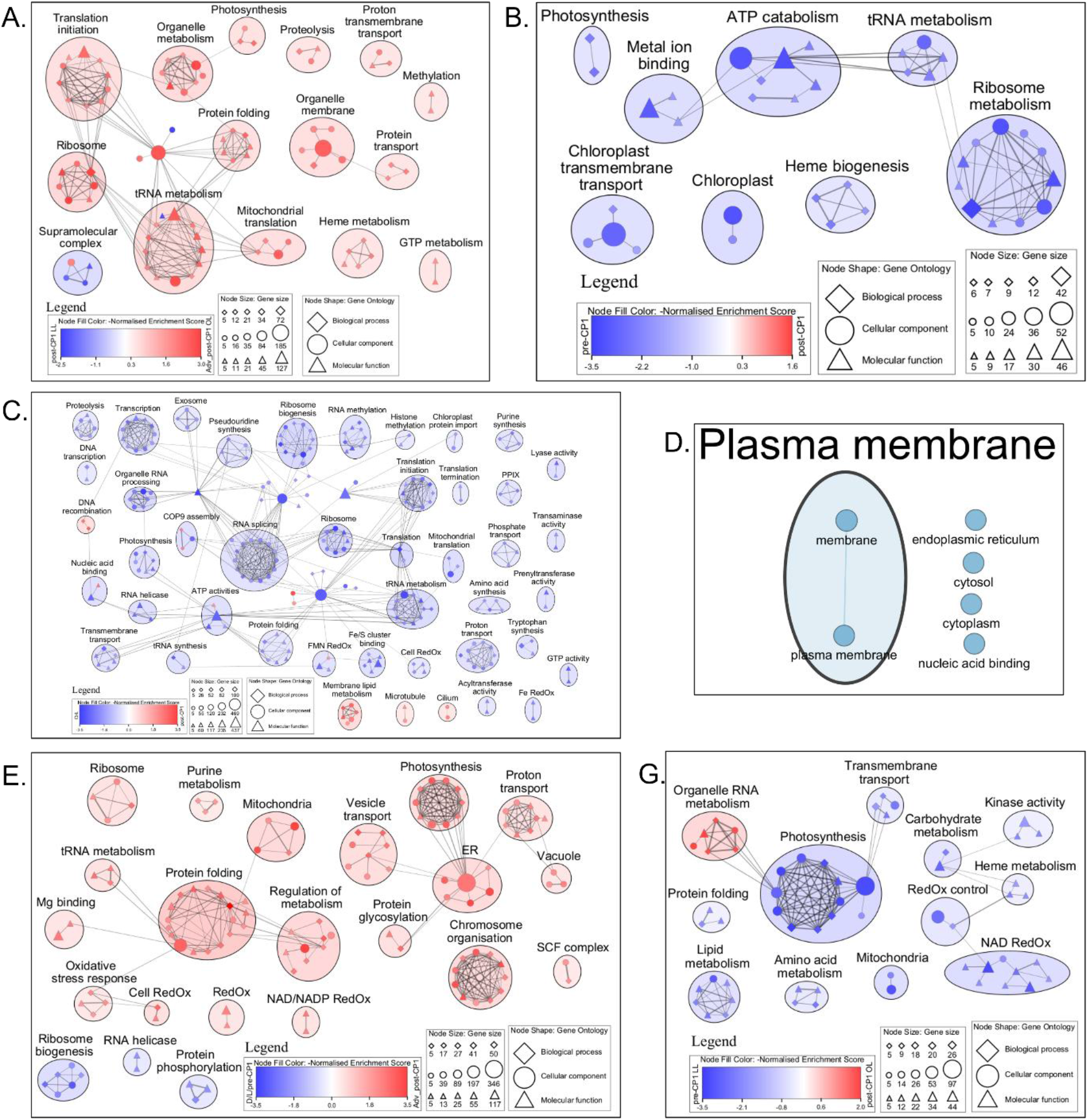
Supplemental Cytoscape figures. The figure was made in Cytoscape from GSEA results using a p-value<0.01, q-value<0.05 and a similarity cutoff Jaccard+overlap coefficient of 0.375. Nodes represent GO terms and clustered together. The single nodes and the nodes’ names have been removed for ease of reading and the details are in supplementary table 2. The symbols represent the GO categories with diamond as biological process; cercle as cellular component and triangle as molecular function. The size of the symbols represents the number of genes in the node. **A.** Network landscape of the GO terms differently enriched between post-CP1 in LL and Adv_post-CP1 in OL. The red color represent Adv_post-CP1 in OL, blue represents post-CP1 in LL. **B.** Network landscape of the GO terms differently enriched between pre-CP1 in LL and post-CP1 in LL. The red color represent post-CP1 in LL, blue represents pre-CP1 in LL. **C.** Network landscape of the GO terms differently enriched between D/L in LL and post-CP1 in LL. The red color represent post-CP1 in LL, blue represents D/L in LL. **D.** Network landscape of the GO terms differently enriched between post-CP1 in OL and Adv_post-CP1 in OL. The blue colour represents Adv_post-CP1 in OL. **E.** Network landscape of the GO terms differently enriched between D/L/pre-CP1 in OL and Adv_post-CP1 in OL. The red color represent Adv_post-CP1 in OL, blue represents D/L/pre-CP1 in OL. **F.** Network landscape of the GO terms differently enriched between pre-CP1 in LL and post-CP1 in OL. The red color represent post-CP1 in OL, blue represents pre-CP1 in LL.

## REFERENCES

1. Khan, M. I., Shin, J. H. & Kim, J. D. The promising future of microalgae: current status, challenges, and optimization of a sustainable and renewable industry for biofuels, feed, and other products. Microbial Cell Factories 2018 17:1 17, 36- (2018).

2. Barsanti, L. et al. Oddities and Curiosities in the Algal World. NATO Security through Science Series C: Environmental Security 353–391 (2008) doi:10.1007/978-1-4020-8480-5_17.

3. Thoré, E. S. J., Muylaert, K., Bertram, M. G. & Brodin, T. Microalgae. Current Biology 33, R91–R95 (2023).

4. Kholssi, R., Lougraimzi, H. & Moreno-Garrido, I. Influence of salinity and temperature on the growth, productivity, photosynthetic activity and intracellular ROS of two marine microalgae and cyanobacteria. Mar. Environ. Res. 186, 105932 (2023).

5. El Shafay, S. M., Gaber, A., Alsanie, W. F. & Elshobary, M. E. Influence of Nutrient Manipulation on Growth and Biochemical Constituent in Anabaena variabilis and Nostoc muscorum to Enhance Biodiesel Production. Sustainability 2021, Vol. 13, 13, (2021).

6. Feng, Y. et al. Using high CO2 concentrations to culture microalgae for lipid and fatty acid production: Synthesis based on a meta-analysis. Aquaculture 594, 741386 (2025).

7. Tan, J. Sen et al. A review on microalgae cultivation and harvesting, and their biomass extraction processing using ionic liquids. Bioengineered 11, 116 (2020).

8. Stirbet, A., Lazár, D., Guo, Y. & Govindjee, G. Photosynthesis: basics, history and modelling. Ann. Bot. 126, 511 (2019).

9. Vecchi, V., Barera, S., Bassi, R. & Dall’osto, L. Potential and Challenges of Improving Photosynthesis in Algae. Plants 9, 67 (2020).

10. Nelson, N. & Junge, W. Structure and energy transfer in photosystems of oxygenic photosynthesis. Annu. Rev. Biochem. 84, 659–683 (2015).

11. Bialevich, V., Zachleder, V. & Bišová, K. The Effect of Variable Light Source and Light Intensity on the Growth of Three Algal Species. Cells 11, 1293 (2022).

12. Kong, F. et al. Effects of light spectra on growth and compositions of biomass, fatty acids, and pigments in three typical microalgae from different phyla. J. Oceanol. Limnol. 42, 1976–1990 (2024).

13. Zhong, Y., Jin, P. & Cheng, J. J. A comprehensive comparable study of the physiological properties of four microalgal species under different light wavelength conditions. Planta 2018 248:2 248, 489–498 (2018).

14. Sartori, R. B. et al. The Role of Light on the Microalgae Biotechnology: Fundamentals, Technological Approaches, and Sustainability Issues. Recent Pat. Biotechnol. 18, 22–51 (2024).

15. Schulze, P. S. C., Guerra, R., Pereira, H., Schüler, L. M. & Varela, J. C. S. Flashing LEDs for Microalgal Production. Trends Biotechnol. 35, 1088–1101 (2017).

16. Tamburic, B., Zemichael, F. W., Maitland, G. C. & Hellgardt, K. Effect of the Light Regime and Phototrophic Conditions on Growth of the H2-producing Green Alga Chlamydomonas Reinhardtii. Energy Procedia 29, 710–719 (2012).

17. Maltsev, Y., Maltseva, K., Kulikovskiy, M. & Maltseva, S. Influence of Light Conditions on Microalgae Growth and Content of Lipids, Carotenoids, and Fatty Acid Composition. Biology (Basel). 10, (2021).

18. Gururani, M. A., Venkatesh, J. & Tran, L. S. P. Regulation of Photosynthesis during Abiotic Stress-Induced Photoinhibition. Mol. Plant 8, 1304–1320 (2015).

19. Hammesfahr, B. & Kollmar, M. Evolution of the eukaryotic dynactin complex, the activator of cytoplasmic dynein. BMC Evol. Biol. 12, (2012).

20. van Hooff, J. J., Tromer, E., van Wijk, L. M., Snel, B. & Kops, G. J. Evolutionary dynamics of the kinetochore network in eukaryotes as revealed by comparative genomics. EMBO Rep. 18, 1559–1571 (2017).

21. Vleugel, M., Hoogendoorn, E., Snel, B. & Kops, G. J. P. L. Evolution and Function of the Mitotic Checkpoint. Dev. Cell 23, 239–250 (2012).

22. Wang, Z. Cell Cycle Progression and Synchronization: An Overview. Methods in Molecular Biology 2579, 3–23 (2022).

23. Rawat, S. S. & Laxmi, A. Sugar signals pedal the cell cycle! *Front*. Plant Sci. 15, (2024).

24. Morgan, David. The cell cycle : principles of control. 297 (2007).

25. Cross, F. R. & Umen, J. G. The Chlamydomonas cell cycle. Plant J. 82, 370–392 (2015).

26. Tulin, F. & Cross, F. R. A Microbial Avenue to Cell Cycle Control in the Plant Superkingdom. Plant Cell 26, 4019 (2014).

27. Zachleder, V., Bišová, K. & Vítová, M. The Cell Cycle of Microalgae. The Physiology of Microalgae 3–46 (2016) doi:10.1007/978-3-319-24945-2_1.

28. Lien, T. & Knutsen, G. SYNCHRONOUS GROWTH OF CHLAMYDOMONAS REINHARDTII (CHLOROPHYCEAE): A REVIEW OF OPTIMAL CONDITIONS1. J. Phycol. 15, 191–200 (1979).

29. Bišová, K. & Zachleder, V. Cell-cycle regulation in green algae dividing by multiple fission. J. Exp. Bot. 65, 2585–2602 (2014).

30. Ivanov, I. N., Vítová, M. & Bišová, K. Growth and the cell cycle in green algae dividing by multiple fission. Folia Microbiol. (Praha). 64, 663–672 (2019).

31. Zachleder, V. et al. Characterization of Growth and Cell Cycle Events Affected by Light Intensity in the Green Alga Parachlorella kessleri: A New Model for Cell Cycle Research. Biomolecules 2021, Vol. 11, **11**, (2021).

32. Zachleder, V. et al. Supra-Optimal Temperature: An Efficient Approach for Overaccumulation of Starch in the Green Alga Parachlorella kessleri. Cells 2021, Vol. 10, **10**, (2021).

33. Ševčíková, T. et al. Completion of cell division is associated with maximum telomerase activity in naturally synchronized cultures of the green alga Desmodesmus quadricauda. FEBS Lett. 587, 743–748 (2013).

34. Moudříková, Š. et al. Comparing Biochemical and Raman Microscopy Analyses of Starch, Lipids, Polyphosphate, and Guanine Pools during the Cell Cycle of Desmodesmus quadricauda. Cells 2021, Vol. 10, **10**, 1–21 (2021).

35. Vítová, M. et al. Chlamydomonas reinhardtii: duration of its cell cycle and phases at growth rates affected by temperature. Planta 234, 599–608 (2011).

36. Harshkova, D. et al. Diclofenac alters the cell cycle progression of the green alga chlamydomonas reinhardtii. Cells 10, (2021).

37. Zachleder, V., Ivanov, I., Vítová, M. & Bišová, K. Cell Cycle Arrest by Supraoptimal Temperature in the Alga Chlamydomonas reinhardtii. Cells 2019, Vol. 8, **8**, (2019).

38. Tamiya, H., Iwamura, T., Shibata, K., Hase, E. & Nihei, T. Correlation between photosynthesis and light-independent metabolism in the growth of Chlorella. Biochim. Biophys. Acta 12, 23–40 (1953).

39. Berková, E., Doucha, J., Vendlová, J. & Zachleder, V. The coupling of energy metabolismus with synthetic and reproduction processes in cell cycles of Scenedesmus quadricauda. Ann. Rep. Algolog. Stud. for 1970 31–51 (1973).

40. Lien, T. & Knutsen, G. Synchronous cultures of Chlamydomonas reinhardti: Synthesis of repressed and derepressed phosphatase during the life cycle. Biochimica et Biophysica Acta (BBA) - Nucleic Acids and Protein Synthesis 287, 154–163 (1972).

41. Harper, J. D. I., Wu, L., Sakuanrungsirikul, S. & John, P. C. L. Isolation and partial characterization of conditional cell division cycle mutants in Chlamydomonas. Protoplasma 186, 149–162 (1995).

42. Morimura, Y. SYNCHRONOUS CULTURE OF CHLORELLA: I. KINETIC ANALYSIS OF THE LIFE CYCLE OF CHLORELLA ELLIPSOIDEA AS AFFECTED BY CHANGES OF TEMPERATURE AND LIGHT INTENSITY. Plant Cell Physiol. 1, 49–62 (1959).

43. Zachleder, V. & Šetlík, I. Timing of events in overlapping cell reproductive sequences and their mutual interactions in the alga Scenedesmus quadricauda. J. Cell Sci. 97, 631–638 (1990).

44. Zachleder, V. & Van Den Ende, H. Cell cycle events in the green alga Chlamydomonas eugametos and their control by environmental factors. J. Cell Sci. 102, 469–474 (1992).

45. Vítová, M. et al. Chlamydomonas reinhardtii: duration of its cell cycle and phases at growth rates affected by light intensity. Planta 233, 75–86 (2011).

46. John, P. C. Control of the cell division cycle in Chlamydomonas. Microbiological sciences, 1(*4*) 96–101 (1984).

47. Zachleder, V. REGULATION OF GROWTH PROCESSES DURING THE CELL CYCLE OF THE CHLOROCOCCAL ALGA SCENEDESMUS QUADRICAUDA UNDER A DNA REPLICATION BLOCK. J. Phycol. 31, 941–947 (1995).

48. Šetlík, I., Ballin, G., Doucha, J. & Zachleder, V. Macromolecular syntheses and the course of cell cycle events in the chlorococcal alga scenedesmus quadricauda under nutrient starvation: Effect of sulphur starvation. Biol. Plant. 30, 161–169 (1988).

49. Panchy, N. et al. Prevalence, evolution, and cis-regulation of diel transcription in Chlamydomonas reinhardtii. G3: Genes, Genomes, Genetics 4, 2461–2471 (2014).

50. Zones, J. M., Blaby, I. K., Merchant, S. S. & Umen, J. G. High-Resolution Profiling of a Synchronized Diurnal Transcriptome from Chlamydomonas reinhardtii Reveals Continuous Cell and Metabolic Differentiation. Plant Cell 27, 2743 (2015).

51. Øezanka, T. et al. The effect of lanthanides on photosynthesis, growth, and chlorophyll profile of the green alga Desmodesmus quadricauda. Photosynth. Res. 130, 335–346 (2016).

52. Èížková, M., Slavková, M., Vítová, M., Zachleder, V. & Bišová, K. Response of the Green Alga Chlamydomonas reinhardtii to the DNA Damaging Agent Zeocin. Cells 2019, Vol. 8, **8**, (2019).

53. Hlavová, M., Vítová, M. & Bišová, K. Synchronization of Green Algae by Light and Dark Regimes for Cell Cycle and Cell Division Studies. Methods in Molecular Biology 1370, 3– 16 (2016).

54. Kselíková, V., Husarčíková, K., Mojzeš, P., Zachleder, V. & Bišová, K. Cultivation of the microalgae Chlamydomonas reinhardtii and Desmodesmus quadricauda in highly deuterated media: Balancing the light intensity. Front. Bioeng. Biotechnol. 10, 960862 (2022).

55. Mackinney, G. ABSORPTION OF LIGHT BY CHLOROPHYLL SOLUTIONS. Journal of Biological Chemistry 140, 315–322 (1941).

56. Lichtenthaler, H. K. & Wellburn, A. R. Determinations of total carotenoids and chlorophylls a and b of leaf extracts in different solvents. Biochem. Soc. Trans. 11, 591–592 (1983).

57. Kinne, O. et al. Buchbesprechungen. Helgoländer Meeresuntersuchungen 1980 34:2 34, 237–250 (1980).

58. Brányiková, I. et al. Microalgae—novel highly efficient starch producers. Biotechnol. Bioeng. 108, 766–776 (2011).

59. Wanka, F. Die Bestimmung der Nucleinsäuren in Chlorella-Kulturen. Planta 58, 594–609 (1962).

60. Decallonne, J. R. & Weyns, J. C. A shortened procedure of the diphenylamine reaction for the measurement of deoxyribonucleic acid by using light activation. Anal. Biochem. 74, 448–456 (1976).

61. Zachleder & Vilém. Optimization of nucleic acids assay in green and blue-green algae: Extraction procedures and the light-activated diphenylamine reaction for DNA. *Algological Studies/Archiv für Hydrobiologie*, Supplement Volumes 67, 313–328 (1984).

62. Lowry, O. H., Rosebrough, N. J., Farr, A. L. & Randall, R. J. PROTEIN MEASUREMENT WITH THE FOLIN PHENOL REAGENT. Journal of Biological Chemistry 193, 265–275 (1951).

63. Chen, S. Ultrafast one-pass FASTQ data preprocessing, quality control, and deduplication using fastp. iMeta 2, e107 (2023).

64. Craig, R. J. et al. The Chlamydomonas Genome Project, version 6: Reference assemblies for mating-type plus and minus strains reveal extensive structural mutation in the laboratory. Plant Cell 35, 644–672 (2023).

65. Patro, R., Duggal, G., Love, M. I., Irizarry, R. A. & Kingsford, C. Salmon provides fast and bias-aware quantification of transcript expression. Nature Methods 2017 14:4 14, 417–419 (2017).

66. 66. Babraham Bioinformatics - FastQC A Quality Control tool for High Throughput Sequence Data. https://www.bioinformatics.babraham.ac.uk/projects/fastqc/.

67. Soneson, C., Love, M. I. & Robinson, M. D. Differential analyses for RNA-seq: transcript-level estimates improve gene-level inferences. F1000Research *2015 4:*1521 4, 1521 (2016).

68. Chen, Y., Chen, L., Lun, A. T. L., Baldoni, P. L. & Smyth, G. K. edgeR v4: powerful differential analysis of sequencing data with expanded functionality and improved support for small counts and larger datasets. Nucleic Acids Res. 53, (2025).

69. Love, M. I., Huber, W. & Anders, S. Moderated estimation of fold change and dispersion for RNA-seq data with DESeq2. Genome Biol. 15, 550- (2014).

70. Usadel, B. et al. A guide to using MapMan to visualize and compare Omics data in plants: a case study in the crop species, Maize. Plant Cell Environ. 32, 1211–1229 (2009).

71. Subramanian, A. et al. Gene set enrichment analysis: A knowledge-based approach for interpreting genome-wide expression profiles. Proc. Natl. Acad. Sci. U. S. A. 102, 15545– 15550 (2005).

72. Mootha, V. K. et al. PGC-1α-responsive genes involved in oxidative phosphorylation are coordinately downregulated in human diabetes. Nature Genetics 2003 34:3 34, 267–273 (2003).

73. Altenhoff, A. M. et al. OMA orthology in 2021: website overhaul, conserved isoforms, ancestral gene order and more. Nucleic Acids Res. 49, D373–D379 (2021).

74. Shannon, P. et al. Cytoscape: A Software Environment for Integrated Models of Biomolecular Interaction Networks. Genome Res. 13, 2498 (2003).

75. R Core Team. R: A Language and Environment for Statistical Computing. https://www.R-project.org/ (2021).

76. Wickham, H. et al. Welcome to the Tidyverse. J. Open Source Softw. 4, 1686 (2019).

77. Wickham, H. ggplot2. ggplot2 10.1007/978-0-387-98141-3 (2009) doi:10.1007/978-0-387-98141-3.

78. Chen, C. et al. TBtools-II: A “one for all, all for one” bioinformatics platform for biological big-data mining. Mol. Plant 16, 1733–1742 (2023).

79. Liu, D., Vargas-García, C. A., Singh, A. & Umen, J. A cell-based model for size control in the multiple fission alga Chlamydomonas reinhardtii. Current Biology 33, 5215–5224.e5 (2023).

80. Mettler, T. et al. Systems Analysis of the Response of Photosynthesis, Metabolism, and Growth to an Increase in Irradiance in the Photosynthetic Model Organism Chlamydomonas reinhardtii. Plant Cell 26, 2310 (2014).

81. Dupuis, S. et al. Too dim, too bright, and just right: Systems analysis of the Chlamydomonas diurnal program under limiting and excess light. Plant Cell 37, koaf086 (2025).

82. Mei, Q. et al. Regulation of DNA replication-coupled histone gene expression. Oncotarget 8, 95005 (2017).

83. Günesdogan, U., Jäckle, H. & Herzig, A. Histone supply regulates S phase timing and cell cycle progression. Elife 3, e02443 (2014).

84. Bannister, A. J. & Kouzarides, T. Regulation of chromatin by histone modifications. Cell Research 2011 21:3 21, 381–395 (2011).

85. Armstrong, C. & Spencer, S. L. Replication-dependent histone biosynthesis is coupled to cell-cycle commitment. Proc. Natl. Acad. Sci. U. S. A. 118, e2100178118 (2021).

86. Bisova, K., Krylov, D. M. & Umen, J. G. Genome-Wide Annotation and Expression Profiling of Cell Cycle Regulatory Genes in Chlamydomonas reinhardtii. Plant Physiol. 137, 475 (2005).

87. Krzywinski, M. et al. Circos: An information aesthetic for comparative genomics. Genome Res. 19, 1639–1645 (2009).

88. Umen, J. G. & Goodenough, U. W. Control of cell division by a retinoblastoma protein homolog in Chlamydomonas. Genes Dev. 15, 1652–1661 (2001).

89. Heldt, F. S., Tyson, J. J., Cross, F. R. & Novák, B. A Single Light-Responsive Sizer Can Control Multiple-Fission Cycles in Chlamydomonas. Current Biology 30, 634–644.e7 (2020).

90. Voigt, J. & Münzner, P. The Chlamydomonas cell cycle is regulated by a light/dark-responsive cell-cycle switch. Planta 172, 463–472 (1987).

91. Kolberg, L. et al. g:Profiler—interoperable web service for functional enrichment analysis and gene identifier mapping (2023 update). Nucleic Acids Res. 51, W207–W212 (2023).

92. Adams, C. C., Jakovljevic, J., Roman, J., Harnpicharnchai, P. & Woolford, J. L. Saccharomyces cerevisiae nucleolar protein Nop7p is necessary for biogenesis of 60S ribosomal subunits. RNA 8, 150–165 (2002).

93. Quenault, T., Lithgow, T. & Traven, A. PUF proteins: repression, activation and mRNA localization. Trends Cell Biol. 21, 104–112 (2011).

94. Wang, M., Ogé, L., Perez-Garcia, M. D., Hamama, L. & Sakr, S. The PUF Protein Family: Overview on PUF RNA Targets, Biological Functions, and Post Transcriptional Regulation. Int. J. Mol. Sci. 19, 410 (2018).

95. Tam, P. P. C. et al. The Puf family of RNA-binding proteins in plants: Phylogeny, structural modeling, activity and subcellular localization. BMC Plant Biol. 10, 44- (2010).

96. Vuong, T. et al. Metamorphosis of a unicellular green alga in the presence of acetate and a spatially structured three-dimensional environment. New Phytologist 245, 1180–1196 (2025).

97. Grossman-Haham, I. et al. Structure of the radial spoke head and insights into its role in mechanoregulation of ciliary beating. Nature Structural & Molecular Biology 2020 28:1 28, 20–28 (2020).

98. Wang, H. et al. The Global Phosphoproteome of Chlamydomonas reinhardtii Reveals Complex Organellar Phosphorylation in the Flagella and Thylakoid Membrane. Molecular & Cellular Proteomics 13, 2337–2353 (2014).

99. Wingfield, J. L. & Lechtreck, K. F. Chlamydomonas Basal Bodies as Flagella Organizing Centers. Cells 2018, Vol. 7, **7**, (2018).

100. Silflow, C. D. et al. The Vfl1 Protein in Chlamydomonas Localizes in a Rotationally Asymmetric Pattern at the Distal Ends of the Basal Bodies. J. Cell Biol. 153, 63 (2001).

101. Wang, Y., Ren, Y. & Pan, J. Regulation of flagellar assembly and length in Chlamydomonas by LF4, a MAPK-related kinase. The FASEB Journal 33, 6431–6441 (2019).

102. Yang, A., Suh, W. I., Kang, N. K., Lee, B. & Chang, Y. K. MAPK/ERK and JNK pathways regulate lipid synthesis and cell growth of Chlamydomonas reinhardtii under osmotic stress, respectively. Scientific Reports 2018 8:1 8, 13857- (2018).

103. Gomez-Osuna, A., Calatrava, V., Galvan, A., Fernandez, E. & Llamas, A. Identification of the MAPK Cascade and its Relationship with Nitrogen Metabolism in the Green Alga Chlamydomonas reinhardtii. Int. J. Mol. Sci. 21, 3417 (2020).

104. Tam, L. W., Wilson, N. F. & Lefebvre, P. A. A CDK-related kinase regulates the length and assembly of flagella in Chlamydomonas. J. Cell Biol. 176, 819–829 (2007).

105. Li, Y., Liu, D., López-Paz, C., Olson, B. J. S. C. & Umen, J. G. A new class of cyclin dependent kinase in chlamydomonas is required for coupling cell size to cell division. Elife 5, (2016).

106. Depège, N., Bellafiore, S. & Rochaix, J. D. Role of chloroplast protein kinase Stt7 in LHCII phosphorylation and state transition in Chlamydomonas. Science 299, 1572–1575 (2003).

107. Bradley, B. A. & Quarmby, L. M. A NIMA-related kinase, Cnk2p, regulates both flagellar length and cell size in Chlamydomonas. J. Cell Sci. 118, 3317–3326 (2005).

108. Davies, J. P., Yildiz, F. H. & Grossman, A. R. Sac3, an Snf1-like serine/threonine kinase that positively and negatively regulates the responses of Chlamydomonas to sulfur limitation. Plant Cell 11, 1179–1190 (1999).

109. Chamovitz, D. A. Revisiting the COP9 signalosome as a transcriptional regulator. EMBO Rep. 10, 352–358 (2009).

110. Harari-Steinberg, O. & Chamovitz, D. A. The COP9 signalosome: mediating between kinase signaling and protein degradation. Curr. Protein Pept. Sci. 5, 185–189 (2004).

111. Wei, N., Serino, G. & Deng, X. W. The COP9 signalosome: more than a protease. Trends Biochem. Sci. 33, 592–600 (2008).

112. Qin, N., Xu, D., Li, J. & Deng, X. W. COP9 signalosome: Discovery, conservation, activity, and function. J. Integr. Plant Biol. 62, 90–103 (2020).

113. Luo, Q., Zou, X., Wang, C., Li, Y. & Hu, Z. The roles of cullins e3 ubiquitin ligases in the lipid biosynthesis of the green microalgae chlamydomonas reinhardtii. Int. J. Mol. Sci. 22, (2021).

114. Li, X. et al. A genome-wide algal mutant library and functional screen identifies genes required for eukaryotic photosynthesis. Nat. Genet. 51, 627 (2019).

115. Núñez-Delegido, E. et al. Mutations in the plant-conserved uL1m mitochondrial ribosomal protein significantly affect development, growth and abiotic stress tolerance in Arabidopsis thaliana. Plant Growth Regul. 105, 429–448 (2025).

116. Plouviez, M. et al. The proteome of Chlamydomonas reinhardtii during phosphorus depletion and repletion. Algal Res. 71, 103037 (2023).

117. Szyszka-Mroz, B. et al. Cold-Adapted Protein Kinases and Thylakoid Remodeling Impact Energy Distribution in an Antarctic Psychrophile. Plant Physiol. 180, 1291–1309 (2019).

118. Kalra, I. et al. Chlamydomonas sp. UWO 241 Exhibits High Cyclic Electron Flow and Rewired Metabolism under High Salinity. Plant Physiol. 183, 588–601 (2020).

119. Tardif, M. et al. PredAlgo: A New Subcellular Localization Prediction Tool Dedicated to Green Algae. Mol. Biol. Evol. 29, 3625–3639 (2012).

120. Cross, F. R. Regulation of Multiple Fission and Cell-Cycle-Dependent Gene Expression by CDKA1 and the Rb-E2F Pathway in Chlamydomonas. Current Biology 30, 1855–1865.e4 (2020).

121. Bertoni, G. Cell Cycle Regulation by Chlamydomonas Cyclin-Dependent Protein Kinases. Plant Cell 30, 271 (2018).

122. Atkins, K. C. & Cross, F. R. Interregulation of CDKA/CDK1 and the Plant-Specific Cyclin-Dependent Kinase CDKB in Control of the Chlamydomonas Cell Cycle. Plant Cell 30, 429–446 (2018).

123. Rochaix, J. D. Regulation and dynamics of the light-harvesting system. Annu. Rev. Plant Biol. 65, 287–309 (2014).

124. van Lis, R. et al. Hybrid cluster proteins in a photosynthetic microalga. FEBS J. 287, 721– 735 (2020).

125. Mus, F., Dubini, A., Seibert, M., Posewitz, M. C. & Grossman, A. R. Anaerobic Acclimation in Chlamydomonas reinhardtii: ANOXIC GENE EXPRESSION, HYDROGENASE INDUCTION, AND METABOLIC PATHWAYS. Journal of Biological Chemistry 282, 25475–25486 (2007).

126. Olson, A. C. & Carter, C. J. The Involvement of hybrid cluster protein 4, HCP4, in Anaerobic Metabolism in Chlamydomonas reinhardtii. PLoS One 11, e0149816 (2016).

127. Wang, J. Z. & Dehesh, K. ER: the Silk Road of interorganellar communication. Curr. Opin. Plant Biol. 45, 171–177 (2018).

128. Angelos, E., Ruberti, C., Kim, S. J. & Brandizzi, F. Maintaining the Factory: The Roles of the Unfolded Protein Response in Cellular Homeostasis in Plants. Plant J. 90, 671 (2017).

129. Chung, K. P., Frieboese, D., Waltz, F., Engel, B. D. & Bock, R. Identification and characterization of the COPII vesicle-forming GTPase Sar1 in Chlamydomonas. Plant Direct 8, e614 (2024).

130. Kim, Y., Terng, E. L., Riekhof, W. R., Cahoon, E. B. & Cerutti, H. Endoplasmic reticulum acyltransferase with prokaryotic substrate preference contributes to triacylglycerol assembly in Chlamydomonas. Proc. Natl. Acad. Sci. U. S. A. 115, 1652–1657 (2018).

131. Chen, S. et al. Stress on the Endoplasmic Reticulum Impairs the Photosynthetic Efficiency of Chlamydomonas. Int. J. Mol. Sci. 25, (2024).

132. Yamaoka, Y. et al. Identification and functional study of the endoplasmic reticulum stress sensor IRE1 in Chlamydomonas reinhardtii. The Plant Journal 94, 91–104 (2018).

133. Michalak, M. Calreticulin: Endoplasmic reticulum Ca2+ gatekeeper. J. Cell. Mol. Med. 28, e17839 (2023).

134. Mariani, P., Navazio, L. & Zuppini, A. Calreticulin and the Endoplasmic Reticulum in Plant Cell Biology. https://www.ncbi.nlm.nih.gov/books/NBK5995/ (2013).

135. Raffaello, A., Mammucari, C., Gherardi, G. & Rizzuto, R. Calcium at the Center of Cell Signaling: Interplay between Endoplasmic Reticulum, Mitochondria, and Lysosomes. Trends Biochem. Sci. 41, 1035–1049 (2016).

136. Goodenough, U. & Roth, R. Ultrastructure of the Endoplasmic Reticulum in Eukaryotic Microalgae. Journal of Eukaryotic Microbiology 72, e70030 (2025).

137. Suzuki, R., Nishii, I., Okada, S. & Noguchi, T. 3D reconstruction of endoplasmic reticulum in a hydrocarbon-secreting green alga, Botryococcus braunii (Race B). Planta 247, 663– 677 (2018).

138. Chen, J., Doyle, C., Qi, X. & Zheng, H. The Endoplasmic Reticulum: A Social Network in Plant CellsF. J. Integr. Plant Biol. 54, 840–850 (2012).

139. Hochmal, A. K., Schulze, S., Trompelt, K. & Hippler, M. Calcium-dependent regulation of photosynthesis. Biochimica et Biophysica Acta (BBA) - Bioenergetics 1847, 993–1003 (2015).

140. Pivato, M., Grenzi, M., Costa, A. & Ballottari, M. Compartment-specific Ca2+ imaging in the green alga Chlamydomonas reinhardtii reveals high light-induced chloroplast Ca2+ signatures. New Phytologist 240, 258–271 (2023).

141. Pivato, M. & Ballottari, M. Chlamydomonas reinhardtii cellular compartments and their contribution to intracellular calcium signalling. J. Exp. Bot. 72, 5312 (2021).

142. Pilon, M., Römisch, K., Quach, D. & Schekman, R. Sec61p Serves Multiple Roles in Secretory Precursor Binding and Translocation into the Endoplasmic Reticulum Membrane. Mol. Biol. Cell 9, 3455 (1998).

143. Peter, J. et al. Fatty acid export (FAX) proteins contribute to oil production in the green microalga Chlamydomonas reinhardtii. Front. Mol. Biosci. 9, 939834 (2022).

144. Bi, X., Corpina, R. A. & Goldberg, J. Structure of the Sec23/24–Sar1 pre-budding complex of the COPII vesicle coat. Nature 2002 419:6904 419, 271–277 (2002).

145. Strenkert, D. et al. Multiomics resolution of molecular events during a day in the life of Chlamydomonas. Proc. Natl. Acad. Sci. U. S. A. 116, 2374–2383 (2019).

146. Pfeffer, S. et al. Dissecting the molecular organization of the translocon-associated protein complex. Nat. Commun. 8, (2017).

147. Singh, R., et al. From Light to Acetate: How Trophic Conditions Shape Growth and Cell Cycle Progression in Chlamydomonas reinhardtii. bioRxiv 2026.03.29.715089 (2026) doi:10.64898/2026.03.29.715089.

148. Hwang, S., Kawazoe, R. & Herrin, D. L. Transcription of tufA and other chloroplast-encoded genes is controlled by a circadian clock in Chlamydomonas. Proc. Natl. Acad. Sci. U. S. A. 93, 996–1000 (1996).

149. Breidenbach, E., Leu, S., Michaels, A. & Boschetti, A. Synthesis of EF-Tu and distribution of its mRNA between stroma and thylakoids during the cell cycle of Chlamydomonas reinhardii. Biochimica et Biophysica Acta (BBA) - Gene Structure and Expression 1048, 209–216 (1990).

150. Zicker, A. A., Kadakia, C. S. & Herrin, D. L. Distinct roles for the 5’ and 3’ untranslated regions in the degradation and accumulation of chloroplast tufA mRNA: identification of an early intermediate in the in vivo degradation pathway. Plant Mol. Biol. 63, 689–702 (2007).

151. Chevalier, B., Turmel, M., Lemieux, C., Monnat, R. J. & Stoddard, B. L. Flexible DNA Target Site Recognition by Divergent Homing Endonuclease Isoschizomers I-CreI and I-MsoI. J. Mol. Biol. 329, 253–269 (2003).

152. Jurica, M. S., Monnat, R. J. & Stoddard, B. L. DNA Recognition and Cleavage by the LAGLIDADG Homing Endonuclease I-Cre I. Mol. Cell 2, 469–476 (1998).

153. Rosen, L. E. et al. Homing endonuclease I-CreI derivatives with novel DNA target specificities. Nucleic Acids Res. 34, 4791–4800 (2006).

154. Monteiro, L. de F. R., Giraldi, L. A. & Winck, F. V. From Feasting to Fasting: The Arginine Pathway as a Metabolic Switch in Nitrogen-Deprived Chlamydomonas reinhardtii. Cells 12, (2023).

155. Gargouri, M. et al. Identification of regulatory network hubs that control lipid metabolism in Chlamydomonas reinhardtii. J. Exp. Bot. 66, 4551–4566 (2015).

156. Valledor, L., Furuhashi, T., Recuenco-Muñoz, L., Wienkoop, S. & Weckwerth, W. System-level network analysis of nitrogen starvation and recovery in Chlamydomonas reinhardtii reveals potential new targets for increased lipid accumulation. Biotechnol. Biofuels 7, 171- (2014).

157. Schmollinger, S. et al. Nitrogen-Sparing Mechanisms in Chlamydomonas Affect the Transcriptome, the Proteome, and Photosynthetic Metabolism. Plant Cell 26, 1410–1435 (2014).

158. Oldenhof, H., Zachleder, V. & Van Den Ende, H. The cell cycle of Chlamydomonas reinhardtii: The role of the commitment point. Folia Microbiol. (Praha). 52, 53–60 (2007).

159. Liu, D., Lopez-Paz, C., Li, Y., Zhuang, X. & Umen, J. Subscaling of a cytosolic RNA binding protein governs cell size homeostasis in the multiple fission alga Chlamydomonas. PLoS Genet. 20, e1010503 (2024).

160. Schmoller, K. M., Turner, J. J., Kõivomägi, M. & Skotheim, J. M. Dilution of the cell cycle inhibitor Whi5 controls budding yeast cell size. Nature 526, 268 (2015).

161. Zatulovskiy, E., Zhang, S., Berenson, D. F., Topacio, B. R. & Skotheim, J. M. Cell growth dilutes the cell cycle inhibitor Rb to trigger cell division. Science 369, 466 (2020).

162. Swaffer, M. P. et al. Transcriptional and chromatin-based partitioning mechanisms uncouple protein scaling from cell size. Mol. Cell 81, S1097–2765(21)00836–4 (2021).

163. Kofler, L. et al. The novel ribosome biogenesis inhibitor usnic acid blocks nucleolar pre-60S maturation. Nature Communications 2024 15:1 15, 7511- (2024).

164. Goodenough, U. W. & Levine, R. P. THE EFFECTS OF INHIBITORS OF RNA AND PROTEIN SYNTHESIS ON THE RECOVERY OF CHLOROPLAST RIBOSOMES, MEMBRANE ORGANIZATION, AND PHOTOSYNTHETIC ELECTRON TRANSPORT IN THE ac-20 STRAIN OF CHLAMYDOMONAS REINHARDI. Journal of Cell Biology 50, 50–62 (1971).

165. Levin, G. In darkness, remember the light: Chlamydomonas retains low- and high-light-induced acclimatory phenotypes in the dark. Plant Cell 37, koaf130 (2025).

166. Redekop, P. et al. Transcriptional regulation of photoprotection in dark-to-light transition— More than just a matter of excess light energy. Sci. Adv. 8, eabn1832 (2022).

167. Liu, X. et al. Hidden diversity: Transcriptomic and photosynthetic variation among common ‘wild type’ Chlamydomonas strains. The Plant Journal 124, e70615 (2025).

